# Parabrachial CGRP Neurons Regulate Opioid Reinforcement

**DOI:** 10.64898/2026.03.18.712659

**Authors:** Lauren L. Bystrom, Alexander V. Margetts, Nicole M. Kujas, Florence M Bourgain-Guglielmetti, Elizabeth P Marinov, Luis M. Tuesta

**Affiliations:** Department of Psychiatry and Behavioral Sciences; Center for Therapeutic Innovation; Sylvester Comprehensive Cancer Center – Cancer Epigenetics Program; Medical Scientist Training Program University of Miami Miller School of Medicine, Miami, FL 33136

## Abstract

Opioid use disorder (OUD) is a chronic, relapsing disease driven by the reinforcing properties of opioids and perpetuated by avoidance of the negative affective states associated with the absence of the drug. Most available OUD treatments directly engage the µ-opioid receptor and may induce side effects that can compromise their therapeutic efficacy, thus underscoring the need for novel therapeutic alternatives. Calcitonin gene-related peptide (CGRP) is produced by a small population of neurons in the parabrachial nucleus (PBN) that has been shown to modulate itch, pain, as well as appetitive behaviors. Using a cell-specific nuclear labeling approach coupled with RNA-sequencing, we generated a baseline transcriptome of CGRP^PBN^ neurons and confirmed expression of multiple genes associated with behavioral responses to appetitive stimuli, as well as enrichment of the µ-opioid receptor, suggesting that CGRP^PBN^ neuron function may be sensitive to the presence of opioids. Indeed, cFos immunostaining showed that CGRP^PBN^ neuron activity increases during early morphine abstinence and reduces gradually over 48 hours. Given the inhibitory effects of opioids on CGRP^PBN^ neuron activity, we next tested whether these neurons could regulate opioid reinforcement. Using a mouse model of morphine intravenous self-administration, we found that chemogenetic inhibition of CGRP^PBN^ neurons significantly reduced the number of morphine rewards earned in both single-dose and dose-response tests but did not affect context-induced morphine seeking after 21 days of abstinence. These results suggest that CGRP^PBN^ neurons are sensitive to opioid administration and can regulate appetitive behaviors such as morphine-taking. Considering that CGRP signaling is regulated by opioid administration, molecular targets that regulate CGRP neurotransmission without direct μ-opioid receptor engagement may therefore serve as novel therapeutic avenues for the treatment of OUD.

**Graphical Abstract:** 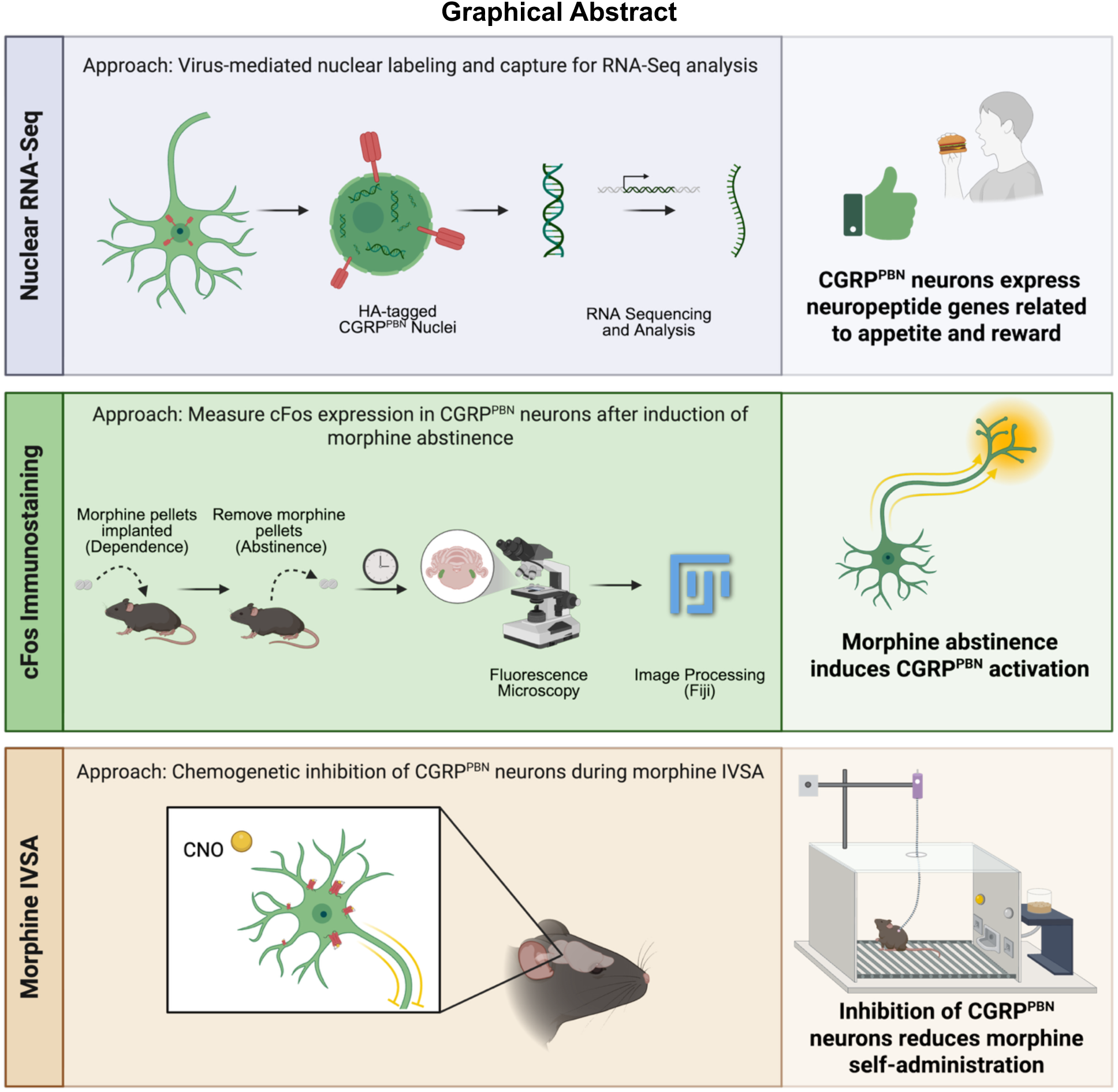

## Introduction

Opioid use disorder (OUD) is a relapsing, remitting disease with significant public health implications. Indeed, the number of opioid-related overdose deaths has increased over the past 40 years, reaching its peak in 2023.^1–3^ While effective therapeutics exist for acute overdose events, therapeutics for cessation and prevention of opioid relapse have been met with limited success.^4,5^ This lack of success is in part due to the mechanism of action of current medication-assisted treatments (MATs), in which drugs such as methadone and buprenorphine directly engage the µ-opioid receptor (µOR), increasing the incidence of negative effects associated with opioid tolerance and withdrawal that can compromise adherence and clinical efficacy.^6–9^ As such, there is a critical unmet need to identify novel therapeutic targets for OUD that function without directly engaging µORs.

Calcitonin Gene-Related Peptide (CGRP) is a neuropeptide produced by a small population of neurons in the lateral external parabrachial nucleus (PBN). CGRP^PBN^ neurons co-release CGRP and glutamate onto brain regions important for motivation, reward, and emotional regulation, including the central amygdala (CeA), ventral tegmental area (VTA), bed nucleus of the stria terminalis (BNST), nucleus accumbens (NAc), and hypothalamus.^10–15^ While peripheral CGRP has important functions, including in migraine pathogenesis, in the central nervous system, CGRP signaling has been studied in relation to pain, itch, taste, sensing of dangerous stimuli, and regulation of appetitive behaviors.^13,16–18^

Preclinical rodent models have previously established that CGRP^PBN^ neurons promote satiety when activated and elicit appetitive signals when inhibited.^13,19–21^ These actions have been shown to be mediated in a CGRP-dependent manner via glucagon-like peptide-1 (GLP-1) projections from the nucleus of the solitary tract (NTS).^19–24^ In addition to its role in appetite regulation, GLP-1 agonism has also been shown to attenuate nicotine, opioid, cocaine, and alcohol taking.^25–29^ Given the behaviorally relevant projections from GLP-1 neurons of the NTS to CGRP^PBN^ neurons and the robust µOR expression within the PBN,^30–32^ it stands to reason that these pathways may also play a role in behavioral responses to opioids. Indeed, the lateral external PBN has been shown to differentially modulate behavior in response to µOR agonists and antagonists.^33–37^ Electrical stimulation of the lateral external PBN elicited both conditioned place preference and aversion responses, depending on the anatomical placement of the electrode. Importantly, both conditioned responses were blocked by application of the µOR antagonist, naloxone.^33–36^ In acute brain slice studies, the lateral external PBN differentially exhibited long-term potentiation or depression to high or low concentrations, respectively, of the synthetic µOR agonist DAMGO.^37^ Additionally, genetic ablation studies in mice have shown some reduction in somatic signs of precipitated morphine withdrawal in ⍺CGRP-deficient mice.^31,38^ Together, accumulating evidence suggests that the PBN has an intrinsic threshold for opioidergic signaling, allowing it to exhibit distinct functional responses at various levels of activation or inactivation. However, the precise role that CGRP^PBN^ neurons may play in regulating behavioral responses to opioids remains poorly understood.

In this study, we first sought to establish the transcriptional profile of CGRP^PBN^ neurons by utilizing a virus-based nuclear labeling and capture technique, followed by RNA-sequencing (RNA-Seq). We then pruned the transcriptome of CGRP^PBN^ neurons by comparing it to those of neighboring cells and other previously published neuron subtypes, with the objective of generating a list of CGRP^PBN^-enriched genes. Next, we established that CGRP^PBN^ neurons are sensitive to opioid abstinence, as measured by cFos immunofluorescence. Lastly, we employed chemogenetic approaches in a model of morphine intravenous self-administration (IVSA) in mice to determine the functional role of CGRP^PBN^ neurons in morphine reinforcement and morphine seeking. These results suggest that CGRP^PBN^ neurons act as regulators of opioid reinforcement, and as such, may represent a novel therapeutic target for the treatment of OUD.

## Methods

### Animals

Homozygous Calca^tm1.1(Cre/EGFP)Rpa^ mice (C57Bl/6 background)(Jackson Laboratories, 033168)^19^ were bred with C57Bl/6J wildtype mice (Jackson Laboratories, 000664) to produce male and female Calca^Cre+/-^ offspring. Male and female Calca^Cre+/-^ animals were used for all experiments and were 8-12 weeks old at the start of each experiment. Animals were housed in the University of Miami Miller School of Medicine’s animal facilities. Mice were maintained on a 12:12 h light/dark cycle and were group-housed with three to five animals per cage. Animals were provided food and water *ad-libitum* for all experiments other than morphine IVSA. In those experiments, animals had free access to food and water until 2 days before the start of behavioral testing, at which time they were food-restricted to 3g of food per mouse per day, with weights taken daily to ensure animals did not fall below 80% of their baseline body weight. All animals were maintained according to National Institutes of Health (NIH) guidelines and Association for Assessment and Accreditation of Laboratory Animal Care (AAALAC) accredited facilities. All experimental protocols were approved by the Institutional Animal Care and Use Committee (IACUC) at the University of Miami Miller School of Medicine.

### Stereotaxic surgery and virus delivery

Anesthesia was induced with a 5% and maintained with a 1-3% isoflurane/ oxygen mixture, and mice were mounted in a stereotaxic frame (Kopf Instruments, Tujunga, CA) at a “flat skull” position. Using aseptic technique, a 5mm longitudinal incision was made on the skin overlying the skull, exposing Bregma, Lambda, and the hindbrain skull surface. Two small circular openings were drilled on the skull to expose the dura mater above the PBN. Bilateral injections (0.375 µL each at 0.2 µL/minute) were made at the coordinates corresponding to the PBN using a 30-gauge syringe (anterior-posterior (AP): -5.0 mm from Bregma; medial-lateral (ML): +/- 1.7 from the midline; dorsal-ventral (DV): -3.2 mm from dura. To ensure appropriate viral dispersion through the PBN, the syringe was left in place for 5 minutes before retracting. For RNAseq experiments, mice received injections of AAV-pEF1a-FLEX-HA-VHH-KASH-WPRE (KASH-HA) as described in Tuesta et al. (2019).^39^ For morphine IVSA, mice received injections of either pAAV-hSyn-DIO-mCherry (Addgene, 50459-AAV5) or pAAV-hSyn-DIO-hM4D(Gi)-mCherry (Addgene, 44362-AAV5).^40^ Bone wax was then used to fill the holes, and the overlying skin was closed with silk sutures. Animals were transferred to a heated incubator to recover from anesthesia and placed in their home cages once fully awake and alert, with access to food and water *ad libitum*. Mice were administered Meloxicam (5mg/kg, s.c.) for analgesia prior to the start of the surgery and 24h postoperatively. The virus was left to incubate for at least 2 weeks before starting behavioral or nuclear isolation procedures.

### Tissue dissection, brain perfusion, and fixation

For RNA-seq, mice were deeply anesthetized using isoflurane, and brains were removed following rapid decapitation. Using a brain block, brains were dissected coronally to reveal the PBN. Bilateral PBN tissue was dissected and combined and transported in Hibernate A Medium (Gibco, A1247501) on ice before dissociation. For histological analyses from all experiments, mice were deeply anesthetized with isoflurane and perfused through the ascending aorta with 1X phosphate buffer saline (PBS; pH 7.4, ThermoFisher, 10010023) plus heparin (15,000 USP units/L; Hepalink, 81952-0112-10), followed by fixation with 4% paraformaldehyde (PFA; Sigma Aldrich, 1002543951) in PBS. Brains were collected, postfixed overnight in 4% paraformaldehyde, then transferred to 30% sucrose in PBS. All brains were cut into 35 µm coronal sections on a cryostat (Leica CM1900), and floating sections were placed into 12-well plates containing PBS with 0.05% sodium azide (Sigma-Aldrich, 1003124924) at 4°C until processing for immunohistochemistry.

### CGRP nuclear isolation, immunolabeling, and fluorescence-activated nuclear sorting (FANS)

All steps in the nuclear isolation process and nuclear immunostaining were performed on ice, and all tubes were pre-chilled. PBN samples were transferred into fresh tubes with Nuclei Extraction Buffer (Miltenyi Biotec, 130-128-024) and were mechanically and enzymatically dissociated on the gentleMACS Octo Dissociator (Miltenyi Biotec, 130-096-427), according to the manufacturer’s instructions. Suspensions were filtered through 100 µm (Miltenyi Biotec, 130-098-463) and 30 µm (Miltenyi Biotec, 130-098-458) filters after repeated centrifugations at 300xg for 5 minutes at 4°C. This nuclear isolate was then resuspended in Wash Buffer (2.5 mM MgCl_2_, 1% BSA, 0.2 U/μL RNAsin in PBS) and incubated for 30 minutes at 4°C before centrifugation at 1000xg for 5 minutes at 4°C. Supernatant was then discarded, and nuclei were resuspended in Wash Buffer with Alexa Fluor 647 Conjugated HA-Tag (6E2) Mouse mAb (1:50, Cell Signaling Technology, 3444S) for 1 hour at 4°C on a tube rotator. After antibody incubation, 64 µL NucBlue Fixed Cell ReadyProbes Reagent (DAPI; Invitrogen, R37606) in 500 µL Wash Buffer was added to nuclei for 20 minutes at 4°C. The immunostained nuclei were then transferred into filtered FACS tubes and sorted with a CytoFLEX SRT (Beckman Coulter) equipped with 15 fluorescent parameters and 2 light scatter parameters at the Flow Cytometry Shared Resource (FCSR) of the Sylvester Comprehensive Cancer Center at the University of Miami. Samples were sorted by nuclear size, physical complexity, positive DAPI signal, and positive AF647 signal. The resulting isolated populations were DAPI^+^/HA^-^ (non-CGRP PBN cells) or DAPI^+^/HA^+^ (CGRP^PBN^ neurons).

### Fluorescence immunolabeling

Free-floating brain sections were processed for fluorescent immunostaining of HA, GFP (CGRP), and cFos. Sections were rinsed in PBN then blocked for 1 hour in Blocking Buffer (10% normal goat serum (Jackson ImmunoResearch; 005-000-079), 0.5% Triton X-100 (Sigma, T8787), and PBS). Sections were then incubated overnight at 4°C in primary antibody diluted in Blocking Buffer. The following primary antibodies were used: Chicken anti-GFP (1:500; Abcam, Ab-13970), Rabbit anti-cFos (1:1000, Cell Signaling Technology, 2250), and Rabbit anti-HA (1:800, Cell Signaling Technology, 3724S). On day 2, sections were washed in PBS three times for 5 minutes each, then incubated with secondary antibody in PBS with 2% normal goat serum for 2 hours at room temperature in the dark. The following secondary antibodies were used: Alexa Fluor 488 Goat Anti-Chicken (1:800, Invitrogen, A11039) and Alexa Fluor 546 Goat Anti-Rabbit (1:800, Invitrogen, A-11035). Following secondary antibody incubation, sections were rinsed in PBS three times for 5 minutes each, mounted on slides with VECTASHIELD Antifade Mounting Medium with DAPI (Vector Laboratories, H-1200–10), and coverslipped. All fluorescent images were taken on an ECHO Revolve Microscope. All antibodies used have been previously validated for the intended applications per the manufacturer, and controls were included to verify background staining by processing the secondary antibodies alone. Only brightness and/or contrast levels were adjusted post-image acquisition and applied to the entire image.

### RNA-Seq and preprocessing

FANS sorted nuclei were directly added to RLT plus buffer (Qiagen, 1053393) for extraction and purification of total RNA according to the manufacturer’s instructions using the Qiagen AllPrep DNA/RNA Mini Kit (Qiagen, 80204). RNA input was normalized across samples, and RNA-Seq libraries were then prepared using NEBNext Single Cell/Low Input RNA Library Prep Kit for Illumina (New England Biolabs, E6420L) according to the manufacturer’s instructions. Sequencing was performed on a Novaseq 6000 (Illumina) or NovaSeq X plus (Illumina) sequencer (paired-end, 150 bp), targeting 30 million reads per sample, by the University of Miami Center John P. Hussman Institute for Human Genomics sequencing core facility and by Azenta Life Sciences (Burlington, MA), respectively. All RNA-Seq data used in this study (produced by our lab or from public datasets (GEO: GSE106956 and GSE63137^39,41^) were mapped to the mm10 genome. Prior to mapping, raw RNA-Seq datasets were trimmed using Trimgalore (v.0.6.7)^42^ and cutadapt (v.1.18).^43^ Illumina sequence adaptors were then removed, and leading and tailing low-quality base-pairs (less than 3) were trimmed. Next, either single- or paired-end reads were mapped to the mm10 mouse genome using STAR (c2.7.10a)^44^ with the following parameters: – outSAMtype BAM SortedByCoordinate –outSAMunmapped Within –outFilterType BySJout – outSAMattributes NH HI AS NM MD XS –outFilterMultimapNmax 20 – outFilterMismatchNoverLmax 0.3 --quantMode TranscriptomeSAM GeneCounts. The resulting bam files were then passed to StringTie (v.2.1.7)^45^ to assemble sequence alignments into estimated transcript and gene count abundance given the NCBI Gencode GRCm38 (NCBI) transcriptome assembly. Reference genome data can be downloaded from the UCSC Genome Browser using the following links: mm10 genome assembly: https://hgdownload.soe.ucsc.edu/goldenPath/mm10/bigZips/mm10.fa.gz and mm10 annotation: https://hgdownload.soe.ucsc.edu/goldenPath/mm10/bigZips/genes/mm10.ncbiRefSeq.gtf.gz

### Differential gene expression analysis

Prior to differential gene expression analysis, genes with fewer than 50 raw count reads across all samples were excluded to reduce noise. Following filtering, the DESeq2^46^ (v. 1.48.1) package from R/Bioconductor (v. 3.21) was used to detect the differentially expressed genes (DEGs) between purified CGRP^PBN^ neurons and publicly available mDA, excitatory, PV, and VIP samples.^39,41^ Only genes with an adjusted *p-value* (padj) < 0.05 were considered to be significantly differentially expressed. When biological replicates showed large variability, indicating outliers, they were removed from each group. For the publicly available datasets, all available data were used, yielding 2 biological replicates per cell type. For the samples collected in our lab, after removing outliers, 2 males and 2 females were left in both the CGRP^PBN^ neuron and the non-CGRP^PBN^ cell groups. Heatmaps were generated using the R/Bioconductor package pheatmap (v.1.0.12) of regular log-transformed (rlog) normalized counts from merged lists of significant DEGs (padj < 0.05). Filters were applied to DEGs to identify the most significant differences between CGRP^PBN^ neurons and all other cell types, which were defined as “CGRP^PBN^-enriched” genes (*padj* < 0.05, log_2_foldchange (L2FC) ≥ 1.5).

### Functional enrichment analysis

The enrichGO function from the R/Bioconductor clusterProfiler package^47,48^ (v. 4.16.0) was used to perform gene ontology (GO) enrichment analysis. All genes meeting the minimum expression threshold established at the beginning of the differential expression analysis (those with >50 transcripts across samples) were included in a list of background genes provided to GO enrichment analyses, and only GO terms and pathways with padj<0.01 and qvalue <0.05 following false discovery rate correction (FDR) were considered. When possible, Fold Enrichment, a score indicating how much a pathway is enriched in a gene list relative to the background list, was set to 3 to identify pathways most strongly associated with CGRP^PBN^ neuron identity. The associated GO plots were generated using the ggplot2^49^ package (v. 3.5.2), with labels added in Adobe Illustrator for clarity.

### Drugs

For the cFos experiment, morphine dependence was established via morphine pellet (25 mg each; NIDA Drug Supply Program, Research Triangle Park, NC, USA) implantation subcutaneously. All pellets were wrapped in nylon mesh prior to implantation. For IV self-administration experiments, morphine sulfate (NIDA Drug Supply Program, Research Triangle Park, NC, USA) was dissolved in 0.9% sterile saline. Drug concentrations were calculated to administer the following doses: 0.1, 0.3, 1.0, and 3.0 mg/kg/infusion. As infusions had a set volume of 35 µl, individual syringe dilutions were calculated based on animal body weight to ensure delivery of an accurate unit dose. Animals in IVSA experiments also received Clozapine-N-Oxide dihydrochloride (CNO; 7.5 mg/kg I.P; Tocris, 6329) dissolved in 0.9% saline or vehicle (0.9% saline) as described in the “Operant IVSA Training” section below.

### Morphine pellet implantation and removal

Animal anesthesia was induced with a 5% and maintained with a 1-3% isoflurane/ oxygen mixture. Under aseptic conditions, animals received 2x25 mg morphine pellets subcutaneously, three days apart. On day 5, both pellets were removed to induce morphine abstinence. After each surgery, the incision was covered with silver sulfadiazine to prevent chewing from group-housed animals and keep the wound clean. Meloxicam (5mg/kg, s.c.) was given for analgesia prior to the start of the surgery. Tissue was collected at 0, 6, 24, and 48 hours post-pellet removal, as described in the tissue dissection section of the methods.

### cFos quantification

After perfusion, free-floating brain sections of the PBN were co-stained for cFos and GFP and imaged on an ECHO Revolve Microscope (San Diego, CA) at 10x magnification. These images were then imported into FIJI^50^ for quantification. The image was converted from RGB to a composite and split into three channels (Blue = DAPI, Red = cFos, and Green = GFP). Using the green channel (GFP), the region of interest of the lateral external PBN was selected and added to the ROI manager. Using the Measurement function, the ROI’s area was calculated. Then, switching to the red channel (cFos), the total number of positive cells only within the ROI was counted. The total number of cFos-positive cells within the ROI was then normalized to its area. This was then repeated for each hemisection on each slice collected from the PBN.

### Jugular catheterization surgery

Jugular catheterization was performed as previously described by Vilca et. al. (2024).^51^ Catheters were made in-house using a 6.5-cm Silastic tube fitted over a guide cannula (PlasticsOne, Protech International Inc., Boerne, TX, USA), bent at a curved right angle, and encased in dental acrylic resin and silicone. Animal anesthesia was induced with a 5% and maintained with a 1-3% isoflurane/ oxygen mixture. Under aseptic conditions, the catheter tubing was passed subcutaneously from the back of the mouse toward the right jugular vein. A small puncture was made in the jugular vein using a 22-gauge guide needle, through which 1 cm of the catheter tip was inserted. After confirming correct placement via blood withdrawal, the catheter was secured with surgical silk sutures. One additional small 2mm incision was made along the back of the mouse, through which the metal cannula portion of the catheter was externalized, and the side incision was closed with sutures. Animals were transferred to a heated incubator to recover from anesthesia and placed in their home cages once fully awake and alert, with access to food and water *ad libitum*. Mice were administered Meloxicam (5mg/kg, s.c.) for analgesia prior to the start of the surgery and 24 h postoperatively. Catheters were flushed daily with 0.9% saline solution containing heparin (10 USP units/mL) starting 48h postoperatively, and done before and after all morphine self-administration sessions. Animals were allowed 3-5 days to recover from surgery prior to operant training.

### Operant IVSA training

Mice were allowed to self-administer IV infusions of morphine during daily 2-hour sessions. Infusions were delivered through Tygon catheter tubing (Braintree Scientific, MA, USA) into the IV catheter using a variable-speed pump (MedAssociated Inc, Fairfax, VT, USA). Operant chambers (MedAssociates Inc., Fairfax, VT, USA) were equipped with grid floors, left- and right-retractable levers, and a yellow cue light above the active lever. Completion of response criteria on the active lever resulted in the intravenous infusion (35 µL over 3 sec) of morphine along with illumination of the cue light for 20 seconds. The period of time after the infusion during which the cue light remained illuminated was a time-out (TO) period. During these 20 seconds, active lever presses were recorded but did not count towards another infusion. Throughout the session, responses on the inactive lever were recorded but had no scheduled consequences. In all IVSA experiments, animals that failed to achieve stable responding (fewer than 7 average infusions earned per 2-hour session during baseline at the 0.3 mg/kg/infusion dose), demonstrated signs of compromised catheter patency, or did not have appropriate viral expression within CGRP^PBN^ neurons (as determined by IHC), were excluded from analysis.

#### Acquisition

During the acquisition phase, mice were allowed to self-administer 0.1 mg/kg/infusion over 5 days of daily 2-hour sessions. The acquisition phase began using a FR1 (fixed ratio 1) TO20 (time out 20s) schedule of reinforcement, where one press of the active lever resulted in a 3s morphine infusion and a 20s TO period. After 5 rewards were earned, the schedule of reinforcement increased to FR2, and subsequently FR3 after 10 earned rewards. On the following day, mice that reached FR3 began at FR2, and those that reached FR1 or FR2 began at FR1. Mice that achieved FR3 for two consecutive days began the next day at FR3. By day 5, all animals began at FR3. Acquisition parameters were identical across all IVSA experiments.

#### Maintenance

Following 8 consecutive days of Acquisition, mice were allowed to self-administer morphine at FR3TO20 during 4 consecutive daily 2-hr maintenance sessions at a dose of 0.3 mg/kg/infusion. Completion of the response criteria on the active lever during maintenance resulted in the delivery of an intravenous infusion (35 μL over 3 sec) of morphine. The average response across maintenance was used to establish a baseline level of responding to the 0.3 mg/kg/infusion dose, which was used in our exclusion criteria.

#### CNO test

After maintenance, mice received an injection of 0.9% saline (10 mL/kg, I.P.) to habituate to the injection protocol and were allowed to respond to morphine. Infusions earned on this day were not counted into either baseline or infusions earned during the vehicle session during the subsequent CNO test. Following saline habituation, the CNO test was conducted using a cross-over design to control for repeated exposure to morphine. All mice received CNO (7.5 mg/kg I.P.) on one day and saline on the other, with wash-out days in between to ensure no lasting effects of the treatment. On washout days, no CNO or vehicle was administered, and animals were allowed to respond to morphine as normal to ensure no lasting effects of chemogenetic inhibition were observed.

#### Dose Response

After acquisition, mice began the dose-response phase, consisting of daily 2-hour sessions at FR3 with correct response criteria resulting in an infusion of morphine, cue light, and a 20-second time out. The doses of morphine tested were 0.1, 0.3, 1.0, and 3.0 mg/kg/infusion. At each dose, a baseline level of responding was first established for at least 2 days, followed by a test day and a washout day. During test days, animals received either CNO (7.5 mg/kg, i.p.) or vehicle 30 minutes prior to the start of the session. On the washout day, no CNO or vehicle was administered, and animals were allowed to respond to morphine as normal to ensure no lasting effects of chemogenetic inhibition were observed.

### Forced home cage abstinence, and context-induced seeking

Following the final session of dose-response testing, animals in the dose-response experiment underwent a context-induced seeking test, representing a 24hr period of drug abstinence (acute abstinence). During this session, no morphine was available; all other parameters remained the same: 2-hour session at FR3 with correct response criteria resulting in pump noises but no morphine infusion, a cue light, and a 20s TO. Total active lever presses were recorded to establish a baseline level of morphine seeking. After this session, mice underwent 21 days of forced home-cage abstinence, during which no behavior or drug was administered. On day 22, animals underwent another context-induced seeking session (chronic abstinence). Animals received either CNO (7.5 mg/kg, i.p.) or vehicle 30 minutes prior to the start of the session, and all other parameters remained consistent with the acute abstinence session: 2-hour session at FR3 with correct response criteria, a cue light, and a 20s TO with no drug delivered. Total active lever presses were recorded as a measurement of motivation to seek the drug.

### Statistical analysis

Sample sizes for experiments were determined using previously published data and power analyses to detect an effect size of d ≥ 1.7 between experimental groups. Statistical analyses were conducted using the packages specified for RNA-Seq data. More information on the specific analyses run is found in the associated code and methods sections above. All other data were assumed to be normally distributed and analyzed using one-way ANOVA, two-way ANOVA with multiple comparisons, or two-way RM ANOVA with Bonferroni post-hoc testing in GraphPad Prism (La Jolla, CA). All animals were randomly assigned to treatment groups, and experiments were blinded whenever possible. Data are shown as mean ± SEM, and α was set at 0.05.

## Results

### Isolated CGRP^PBN^ neurons are enriched for neuropeptide and neuronal markers

To isolate CGRP^PBN^ neurons, we used a previously established virus-mediated nuclear label and capture technique.^39^ Calca^Cre+/-^ heterozygous mice^19^ received bilateral PBN injections of a Cre-inducible AAV-DIO-KASH-HA (**Supplementary Figure 1A**), encoding a hemagglutinin (HA) tag fused to the nuclear membrane protein KASH. Using the endogenous GFP marker expressed in these Cre^+^ cells, we confirmed that Cre^+^ cells in AAV-DIO-KASH-HA-injected Calca^Cre+/-^ mice expressed the HA tag while Cre^-^ cells did not (**Supplementary Figure 1B-C**), indicating that the KASH-HA construct in this transgenic line demonstrated Cre dependence and appropriate infectivity. In a separate cohort of animals, Calca^Cre+/-^ mice received bilateral injections of AAV-DIO-KASH-HA, and two weeks later, tissue was collected and processed for nuclear RNA-Seq (**Figure 1A**). To confirm that HA^+^ nuclei represented CGRP^PBN^ neurons, we performed differential gene expression analysis on HA^+^ and HA^-^ nuclei. This analysis suggested that the highest source of variance between groups (PC1:81%) was cell type (**Figure 1B**). This was further supported by unsupervised clustering analysis, which showed that gene expression profiles clustered according to the presence of the HA tag (**Figure 1C**). Within each group, there was further unsupervised clustering by sex, suggesting that the transcriptional profile of CGRP^PBN^ neurons may be sexually dimorphic (**Supplementary Figure 2A-D**). Further, significant increases in neuron-specific developmental genes, including ISL1 transcription factor (*Isl1*) (*padj* < 0.0001) and chondrolectin (*Chodl*) (*padj* < 0.0001), were enriched in HA^+^ nuclei (**Figure 1D**), while in HA^-^nuclei, there was an upregulation of glial and oligodendrocyte markers, including fatty acid 2-hydroxylase enzyme (*Fa2h*) (*padj* < 0.0001), kallikrein-related peptidase 6 (*Klk6*) (*padj* < 0.0001), glutamine synthetase (*Glul*) (*padj* < 0.0001), and transmembrane protein 63a (*Tmem63a*) (*padj* < 0.0001) (**Figure 1D**). In addition to these, HA^+^ nuclei were enriched for neuropeptide signaling-related genes, including calcitonin-related polypeptide alpha (*Calca*) (*padj* < 0.0001) (encodes CGRP), urotensin 2b (*Uts2b*) (*padj* < 0.001), galanin (*Gal*) (*padj* < 0.0001), and secretogranin II (*Scg2*) (*padj* < 0.05) (**Figure 1D**). Enrichment of these DEGs suggests that cells in the HA^+^ group may play important neuromodulatory roles. Indeed, gene ontology (GO) analysis revealed that significant DEGs in HA^+^ nuclei are enriched for neuronal development, neuropeptide signaling, and pain and stress pathways (*padj_genes_* < 0.05, *padj_pathways_* < 0.01, *q-value_pathways_* < 0.05, L2FC > 1, Fold Enrichment > 3) (**Figure 1E**). DEGS in HA^-^ nuclei were enriched in non-neuronal pathways and immune-related functions (*padj_genes_* < 0.05, *padj_pathways_* < 0.01, *q-value_pathways_* < 0.05, L2FC < -1, Fold Enrichment > 3) (**Figure 1F**). These results confirm that the HA^+^ population represents CGRP^PBN^ neurons, whereas the glial-enriched HA^-^ population consists of non-CGRP PBN cells.

**Figure 1:**
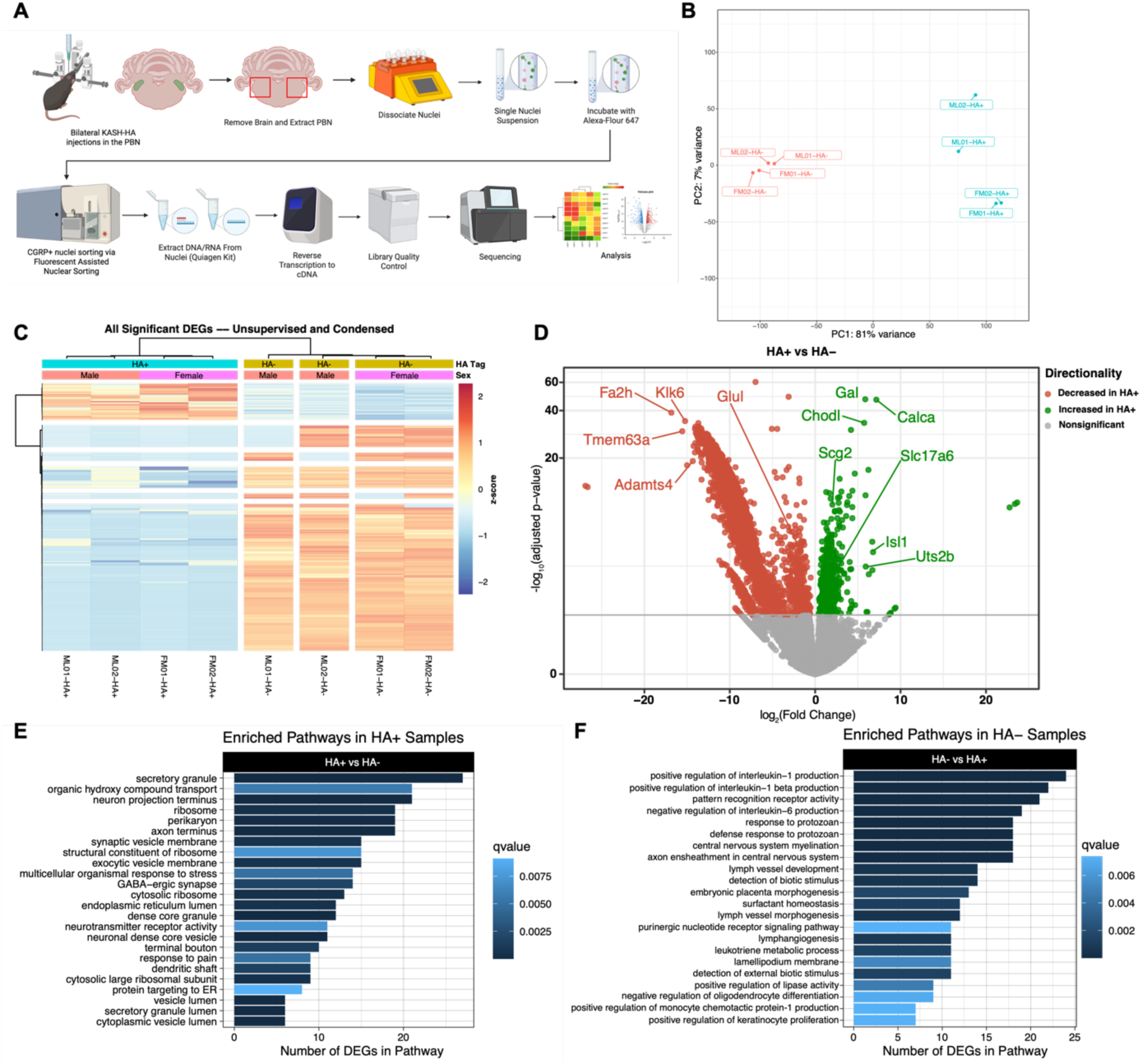
Isolated CGRP^PBN^ nuclei are enriched in neuronal markers and genes related to hormone and neuropeptide signaling, while non-CGRP nuclei express non-neuronal markers. Male and Female Calca^Cre+/-^ mice were bilaterally injected with an AAV-DIO-KASH-HA virus. Two weeks later, the parabrachial nucleus was dissected, and nuclei were isolated using Fluorescence Assisted Nuclear Sorting (FANS). These nuclei were then used for RNAseq analysis. A) Experimental design for the nuclear capture technique and subsequent library preparation for RNAseq. B) Principal Component Analysis of Variance between HA+ nuclei (cyan) and HA- nuclei (salmon) using the top 500 most variable genes. C) Unsupervised clustering heatmap of normalized counts (rlog transformed) of all significantly differentially expressed genes (*padj* < 0.05) between HA+ and HA- nuclei. D) Volcano plot of differentially expressed genes between HA+ and HA-. Green (increased) and red (decreased) circles represent genes that reached significance (*padj* < 0.05). E) Significantly enriched Gene Ontology (GO) pathways in the 609 HA+ enriched genes (*padj_genes_* < 0.05, *padj_pathways_* < 0.01, *q-value_pathways_* < 0.05, L2FC > 1, Fold Enrichment ≥ 3). F) Significantly enriched GO pathways in the 3401 HA- enriched genes (*padj_genes_* < 0.05, *padj_pathways_* < 0.01, *q-value_pathways_* < 0.05, L2FC < -1, Fold Enrichment ≥ 3).

### CGRP^PBN^ identity genes suggest a role in appetite and reward

While the comparison of HA^+^ to HA^-^ nuclei revealed that CGRP^PBN^ neurons are depleted of glial gene expression, many genes expressed in HA^+^ neurons are also expressed in other neuronal subtypes (**Figure 1E**). To identify those genes that are highly enriched specifically in CGRP^PBN^ neurons, we compared the gene expression of this neuronal subtype against publicly available datasets of midbrain dopaminergic (mDA) neurons,^39^ cortical excitatory pyramidal neurons, vasoactive intestinal protein (VIP)- cortical interneurons, and parvalbumin (PV)- cortical interneurons.^41^ Principal Component Analysis (PCA) of these transcriptomes showed that CGRP^PBN^ neurons clustered independently from other neuron subtypes (**Figure 2A**). Next, we generated heatmaps using k-means clustering of all significant DEGs to identify gene groups enriched in each cell type (*padj* < 0.05) (**Figure 2B**). To identify the most significantly upregulated genes in CGRP^PBN^ neurons, L2FC filters were passed and revealed 519 “CGRP^PBN^-enriched genes” (*padj* < 0.05, L2FC ≥ 1.5) (**Figure 2B**). GO analysis indicated that CGRP^PBN^-enriched genes were significantly enriched in pathways related to neuropeptide and neuroendocrine signaling and GABA receptors (*padj_genes_* < 0.05, L2FC > 1.5, *padj_pathways_* < 0.01, *q-value_pathways_* < 0.05, Fold Enrichment ≥ 3) (**Figure 2C**). Further neuropeptides known for their role in appetite and reward, including *Calca* (*padj* < 0.001), *Gal* (*padj* < 0.0001), neurotensin (*Nts*) (*padj* < 0.05), and nesfatin-1 (*Nuc2b*) (*padj* < 0.001), were among the most highly enriched (**Figure 2D**).^13,19,52–57^ Consistent with previous literature, there was also a high mean expression of *Oprm1* in CGRP^PBN^ neurons (mean normalized gene counts: CGRP^PBN^ neurons, 280.75; PBN, 139.5; mDA, 49; Excitatory, 41; PV, 49; VIP, 258.5) (**Supplementary Figure 3A**). Additionally, proenkephalin (*Penk*) (*padj* < 0.05) (endogenous opioid), substance P receptor (*Tacr1*) (*padj* < 0.05) (highly associated with µ-opioid receptor (*Oprm1*)),^30,58,59^ and the nociceptin receptor (*Oprl1*) (*padj* < 0.01) were also included among CGRP^PBN^-enriched genes (**Figure 2D**), highlighting that CGRP^PBN^ neurons may be predisposed to respond to cues related to appetite, reward, and opioid signaling.^11,19,59,60^

**Figure 2:**
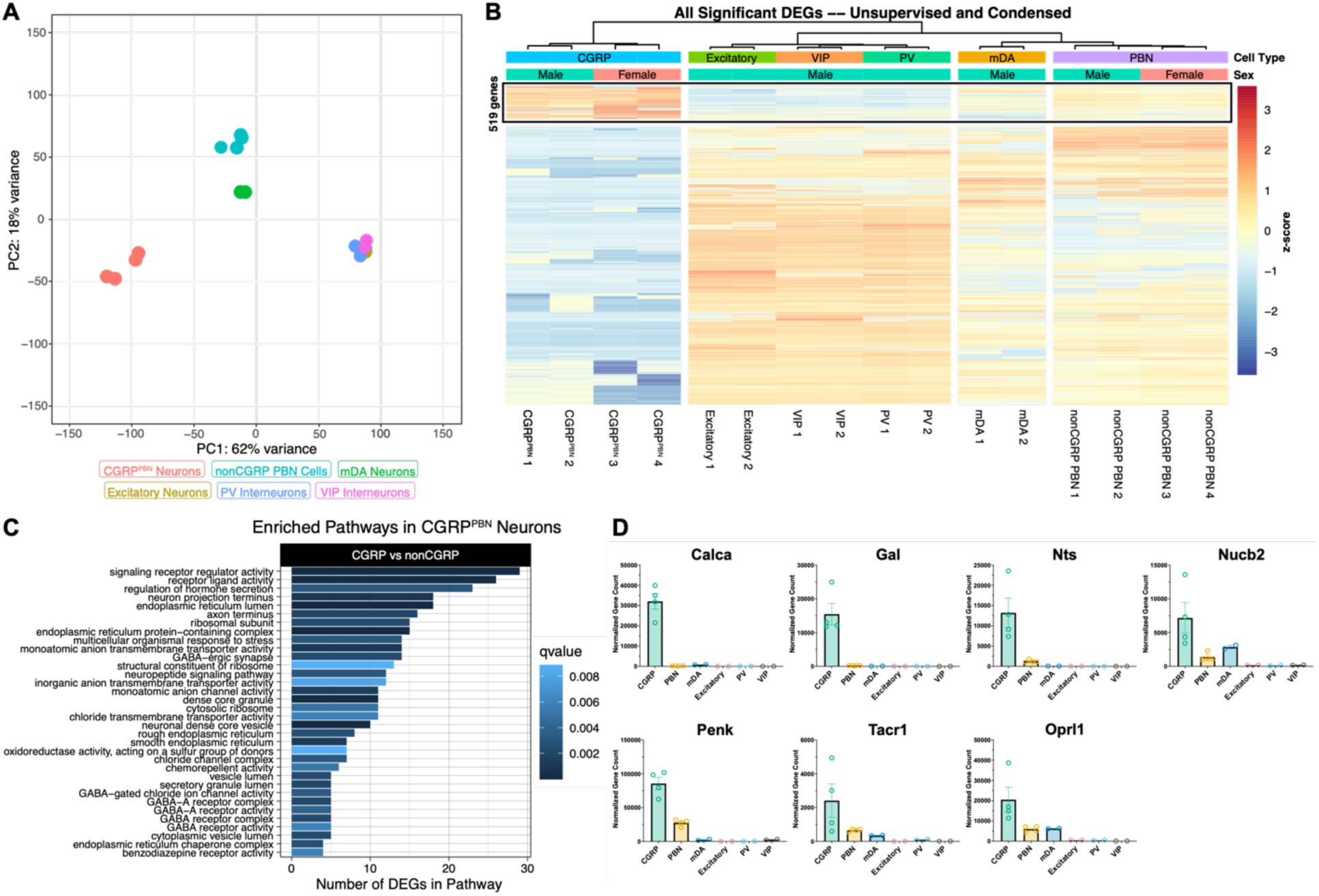
Transcriptome of CGRP^PBN^ neurons is enriched for biological pathways related to neuropeptide processing and signaling, with key identity genes related to addiction and SUDs. Gene expression data from publicly available datasets (GEO: GSE106956 and GSE63137) were processed alongside CGRP^PBN^ and non-CGRP PBN samples and underwent differential gene expression and functional enrichment analyses. **A)** Principal Component Analysis (PCA) using the top 500 genes indicates CGRP^PBN^ neurons (salmon) are transcriptionally distinct from all other brain regions. **B)** Unsupervised clustering heatmap of normalized counts (rlog transformed) of all significantly differentially expressed genes (*padj* < 0.05) between all cell types. The 519 “CGRP^PBN^-enriched” genes (black box) were determined by the most highly significantly upregulated genes in CGRP^PBN^ samples compared to all other cell types (*padj* < 0.05, L2FC ≥ 1.5). **C)** Significantly enriched GO pathways in the CGRP^PBN^-enriched dataset indicate biological pathways most relevant to CGRP^PBN^ neuron identity (*padj_genes_* < 0.05, L2FC ≥ 1.5, *padj_pathways_* < 0.01, *q-value_pathways_* < 0.05, Fold Enrichment ≥ 3). **D)** Normalized gene expression plots highlight important reward and SUD-related genes included in the “CGRP^PBN^-enriched” dataset (*padj* < 0.05, L2FC ≥ 1.5); n=2-4 per cell type; Error bars represent mean ± SEM.

### CGRP^PBN^ neuron activity is sensitive to morphine exposure and abstinence

While previous studies have shown that opioid signaling affects synaptic plasticity and behavior mediated by the PBN, it is still unclear how CGRP^PBN^ neurons respond to opioids and opioid abstinence. ^33–37^ To this end, Calca^Cre+/-^ animals were exposed to 5 consecutive days morphine administration (via subcutaneous pellets), and brains were collected during pellet removal, and at 6, 24, or 48 hours after removal, as illustrated in **Figure 3A**. Sections of the PBN (**Figure 3D**) were then stained and quantified in FIJI. At 0 hours, the average number of cFos^+^ CGRP^PBN^ neurons per mm^2^ was 240.2 ± 18.9 compared to 705.4 ± 25.1 at 6 hours, suggesting an increase in CGRP^PBN^ neuron activation during this time (p < 0.0001) (**Figure 3B-C**). 24 hours after pellet removal, the number of cFos^+^ CGRP^PBN^ neurons per mm^2^ decreased to 558 ± 20.9, and further decreased to 336.5 ± 15.2 by 48 hours (**Figure 3B-C**). The decrease in activation from 6 hours (24 hours: p < 0.0001, 48 hours: p < 0.0001) is consistent with known timelines for acute opioid withdrawal-related behaviors in mice. ^61,62^ Importantly, the number of cFos^+^ CGRP^PBN^ neurons per mm^2^ remained elevated at 48 hours of abstinence compared to 0 hours (p < 0.01) (**Figure 3B-C**). These results suggest that CGRP^PBN^ neurons are sensitive to morphine, as evidenced by the increase in activation 6 hours after pellet removal, and the gradual decrease in activation from 24 to 48 hours.

**Figure 3:**
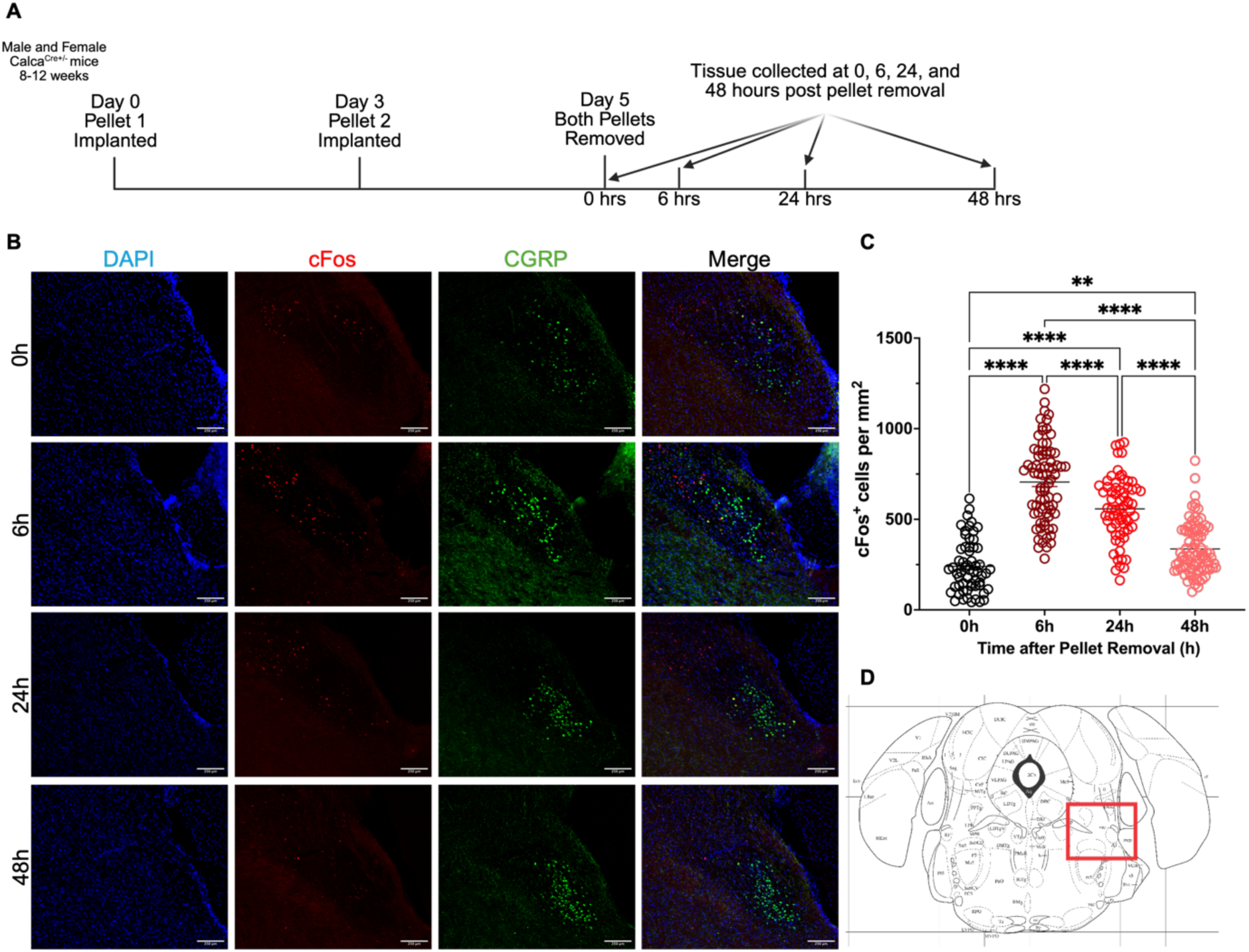
CGRP^PBN^ Neurons are activated by morphine abstinence. Male and Female Calca^Cre+/-^ mice were implanted with 2x25mg morphine pellets three days apart. Both pellets were removed on day 5 to induce morphine abstinence. Tissue was collected at 0, 6, 24, or 48 hours post-pellet removal, then sectioned and stained for cFos. Localization of CGRP^PBN^ neurons was confirmed by GFP staining for the endogenous GFP expressed within Calca^Cre+/-^ animals. Quantification of cFos+ cells within this region of interest was conducted in FIJI. **A)** Experimental Timeline. **B)** Representative IHC images of cFos expression in CGRP^PBN^ neurons from each time point. Images were taken at 10x. Scale bar represents 250 µm. **C)** Quantification of the number of cFos+ cells within the GFP+ region normalized to the area of the ROI. Each point represents one hemisection with 8-18 hemisections included per animal. Ordinary One Way ANOVA: F(3,285) = 104.8, p<0.0001; **p<0.01, ****p<0.0001; N=22: 0h n=5, 6h n=5, 24h n=6, 48h n=6. Error bars represent mean ± SEM. **D)** Schematic Image from Allen Brain Atlas representing the region from which images were taken.

### Chemogenetic inhibition of CGRP^PBN^ neurons reduces morphine taking

Considering that CGRP^PBN^ neurons are sensitive to morphine administration and abstinence, we next asked whether this population played a role in regulating the appetitive aspects of opioid-taking. As described in **Figure 4A**, we used a model of morphine IVSA in which Calca^Cre+/-^ mice received bilateral PBN injections of either a control virus (pAAV-hSyn-DIO-mCherry) or an inhibitory DREADD virus (pAAV-hSyn-DIO-hM4D(Gi)-mCherry). As with the KASH-HA, the hM4Di construct was validated to show Cre-dependence and appropriate infectivity (**Supplementary Figure 4A-C**). Two weeks after viral delivery, animals were implanted with jugular catheters and underwent operant training, quickly establishing and maintaining stable morphine-taking (**Figure 4B**). Chemogenetic inhibition of CGRP^PBN^ neurons by CNO resulted in a significant decrease in the number of morphine infusions earned (9.7 ± 1.4 infusions) compared to vehicle injection (15 ± 1.5 infusions) (Paired t-test, p<0.001) (**Figure 4D**). Importantly, there was no significant difference in the number of infusions earned between CNO- (14.9 ± 3.0 infusions) and vehicle-injected (13.4 ± 2.3 infusions) control (DIO-mCherry) mice (**Figure 4C**), suggesting that the inhibitory effects of CNO on morphine reinforcement are restricted to CGRP^PBN^ neurons.

**Figure 4:**
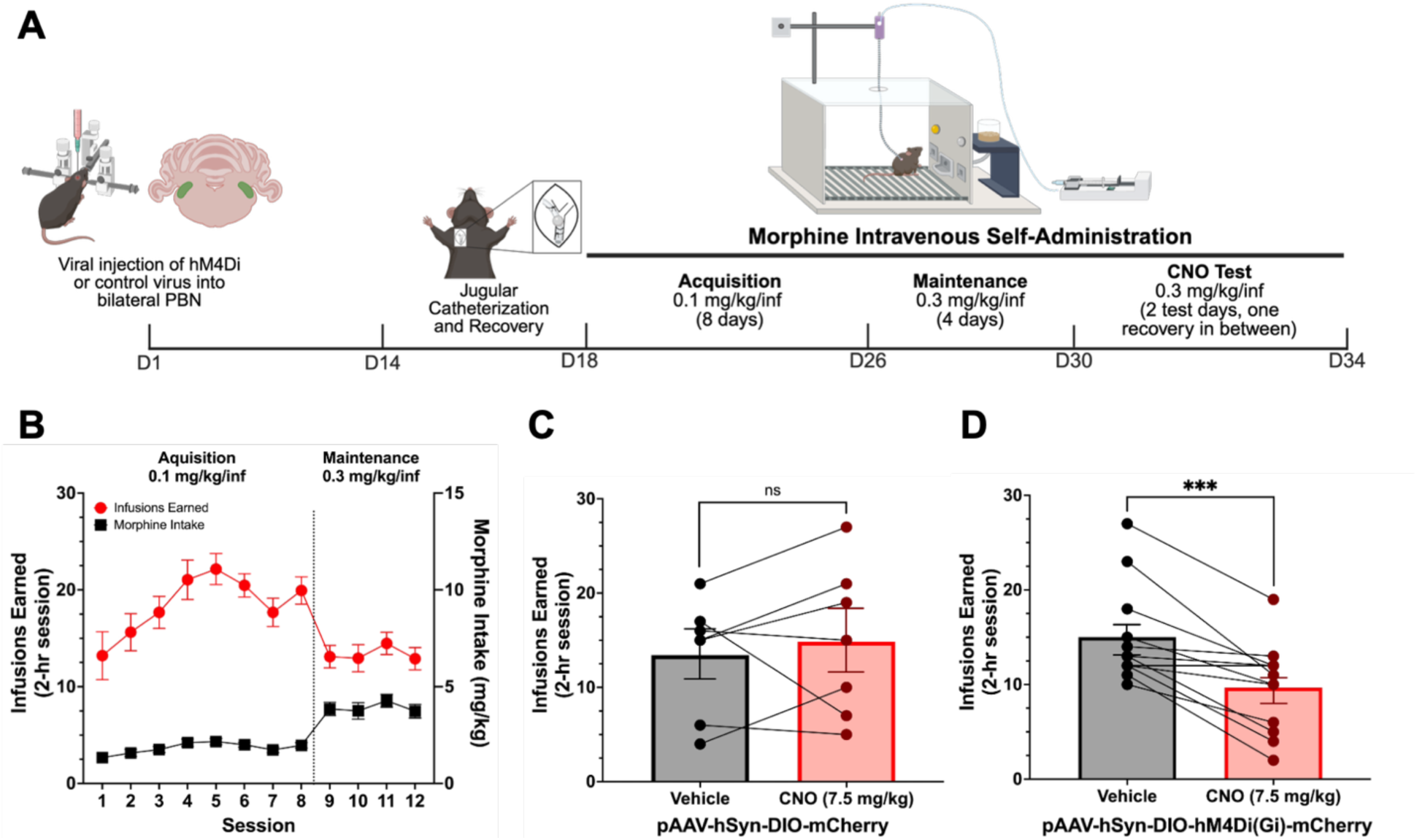
Chemogenetic inhibition of CGRP^PBN^ neurons reduces morphine-taking. Male and female Calca^Cre+/-^ mice received bilateral PBN injections of hM4Di or control virus. Two weeks later, mice underwent jugular catheterization and were trained to self-administer morphine during daily 2-hour sessions at FR3TO20. Mice received either CNO (7.5 mg/kg, I.P) or vehicle 30 minutes prior to session start during test sessions. **A)** Experimental Timeline **B)** Number of infusions earned (red) and total morphine intake (black) during each daily behavioral session across all animals (n=19). **C)** Number of infusions earned during vehicle and CNO sessions in control virus-injected mice (n=7). **D)** Number of infusions earned during vehicle and CNO session in DREADD-injected mice (n=12). Paired t-test (Infusions earned during CNO vs vehicle session, ***p<0.001). **B-D)** Data are represented as mean ± SEM.

### Chemogenetic inhibition of CGRP^PBN^ neurons flattens the dose response curve but does not affect morphine reinstatement

While the data presented in **Figure 4** indicate that CGRP^PBN^ neurons regulate opioid reinforcement, only one morphine dose was tested. Therefore, we next asked whether this effect would be consistent across various morphine doses in a dose-response model. To this end, Calca^Cre+/-^ mice received bilateral PBN injections of pAAV-hSyn-DIO-hM4D(Gi)-mCherry. Two weeks later, animals were implanted with jugular catheters and underwent operant training to receive intravenous infusions of morphine at the following doses: 0.1, 0.3, 1.0, and 3.0 mg/kg/infusion, as described in **Figure 5A**. For each dose tested, mice were allowed to establish a baseline level of responding over two days before test sessions in which CNO (7.5 mg/kg, i.p.) or vehicle was administered. CNO-treated mice decreased the average number of morphine infusions earned compared to baseline at two of the four doses tested (Mixed Effects ANOVA with Bonferroni post-hoc- Effect of CNO-treatment (F(1,18)=5.25, p<0.05)) (**Figure 5B**, **Supplementary Figure 5A**). There was no statistical difference in the number of morphine infusions earned by vehicle-treated animals in test vs baseline sessions at any of the doses tested (**Figure 5C**, **Supplementary Figure 5A**), suggesting that inhibition of CGRP^PBN^ neurons reduces the motivation to consume opioids.

**Figure 5:**
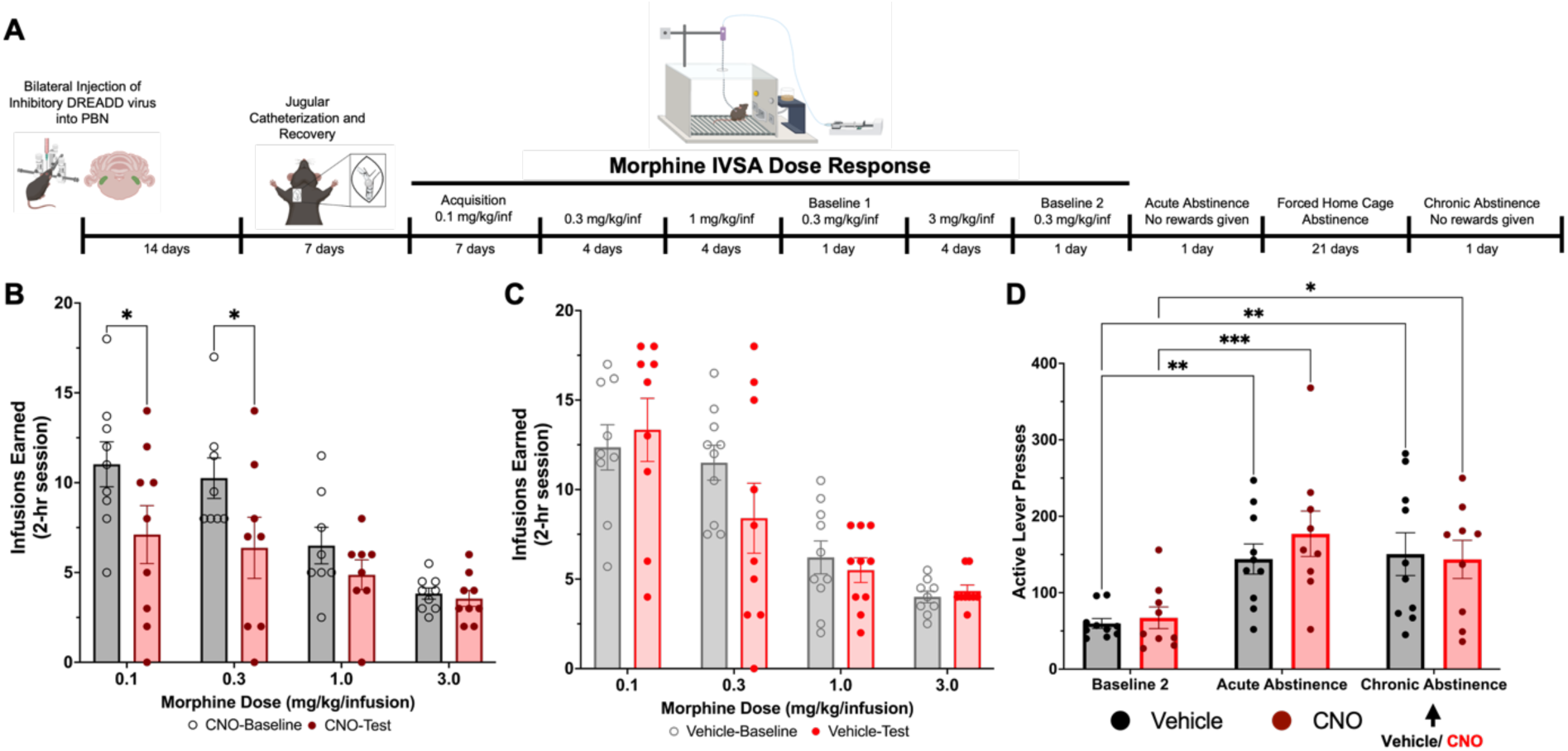
CGRP^PBN^ neurons regulate morphine reinforcement, but not morphine-seeking. Male and female Calca^Cre+/-^ mice were injected with an inhibitory DREADD virus. Two weeks later, mice were implanted with jugular catheters and underwent operant training to respond for morphine intravenously in a dose-response model. During test sessions and Chronic Abstinence, mice received either CNO (7.5 mg/kg, I.P) or vehicle 30 minutes prior to session start. **A)** Experimental Timeline. **B)** Morphine infusions earned by CNO-administered animals in baseline and test sessions. (Mixed Effects ANOVA with Bonferroni post-hoc- Effect of CNO-treatment (F(1,18)=5.25, p<0.05). Baseline is represented in black, while test session is represented in maroon. **C)** Morphine infusions earned by Vehicle-administered animals in baseline and test sessions. Baseline is represented in gray, while test session is represented in red. **D)** Number of Active Lever Presses during last morphine session (Baseline 2) compared to Acute and Chronic Abstinence. CNO or vehicle was only administered before the Chronic Abstinence session, as shown by the black arrow. Two-way RM ANOVA- Effect of Session (F(2,34)=17.26, p<0.0001). Vehicle is represented in black, while CNO-treated animals are represented in red. **B-D)** *p<0.05,**p<0.01,***p<0.001; N=9-11 animals per group per comparison. Error bars represent mean ± SEM.

Twenty-four hours after completing the dose-response phase of the morphine IVSA experiment, mice were subjected to 2 context-induced morphine seeking tests separated by 21 days of forced home cage abstinence, as previous studies have shown this to be an effective model for measuring long-lasting drug-seeking behaviors.^63–65^ During these sessions, correct completion of the response criteria resulted in presentation of a cue light, followed by a 20s TO, but with no drug delivery. No CNO or vehicle was administered during acute abstinence, as this session was used to establish an initial level of morphine-seeking. When compared to the morphine taking session from the final IVSA day (Baseline 2), there was a significant increase in the number of active lever presses from 67.2 to 177.1 in the CNO group and from 59.8 to 144.1 in the vehicle group (Two-way RM ANOVA- Effect of Session (F(2,34) = 17.26, p < 0.0001)(**Figure 5D**), suggesting that both groups of animals exhibit morphine-seeking behavior as early as 24 hours of abstinence. No significant differences were noted in baseline craving between the CNO and vehicle groups (Average active lever press acute abstinence: CNO- 177.1 ± 29.7, Vehicle-144.1 ± 19.7)(**Figure 5D**, **Supplementary Figure 5B**). These results were then compared to those from chronic abstinence (21 days), in which animals received either CNO (7.5 mg/kg, I.P.) or vehicle 30 minutes prior to the start of the session. Comparing to Baseline 2, there was also a significant increase in the number of active lever presses from 67.2 to 143.7 in the CNO group and from 59.8 to 150.5 in the vehicle group (Two-way RM ANOVA- Effect of Session (F(2,34) = 17.26, p < 0.0001)(**Figure 5D**). However, no significant differences were noted in active lever responses between acute and chronic abstinence (Average active lever press acute abstinence: CNO- 177.1 ± 29.7, Vehicle- 144.1 ± 19.7; Average active lever press chronic abstinence: CNO-143.7 ± 25.0, Vehicle- 150.5 ± 28.0)(**Figure 5D**), nor between CNO or vehicle-treated animals in chronic abstinence (**Figures 5D**, **Supplementary Figure 5C**). These results suggest that the morphine-seeking behavior displayed during acute abstinence persists through 21 days of forced home cage abstinence, and that this persistent effect is not dependent on the activity of CGRP^PBN^ neurons.

## Discussion

OUD continues to be an economic and public health crisis, despite a recent decline in opioid-related overdose deaths that can be partly attributed to the efficacy and broad availability of therapeutics for acute overdose events.^1^ Unfortunately, the efficacy of current MATs for opioid cessation and relapse prevention is limited in part due to direct µOR engagement, which can compromise patient adherence owing to complications inherent to opioid-based treatments, such as tolerance, withdrawal, and restricted access in the case of methadone.^4,6,8,9,66–70^ As such, there is currently an unmet need for therapeutic targets that function independently of direct µOR engagement. To this end, we propose that CGRP^PBN^ neurons regulate opioid reinforcement and may present such an alternative.

Single-cell (sc)RNA-Seq studies in the PBN have identified CGRP^PBN^ neurons as a distinct population within this relatively heterogeneous brain region.^11^ While scRNA-Seq is optimal for identifying cell populations based on enrichment of highly expressed genes, its limited sequencing depth is insufficient to generate a transcriptional profile that reliably represents lowly-expressed genes. To this end, we first sought to generate a public dataset that comprehensively characterizes CGRP^PBN^ neurons via nuclear RNA-Seq. Using Calca^Cre+/-^ transgenic mice in combination with a Cre-inducible KASH-HA virus enabled cell type-specific tagging of CGRP^PBN^ nuclei for molecular analysis. While this technique has previously been used for other cell types,^39,71^ it had not yet been established within this brain region. Here, we show that this technique is effective for transcriptional profiling of CGRP^PBN^ neurons (**Supplementary Figure 1B-C**) and enables interrogation of key genes expressed within this neuronal population, providing greater depth of coverage than single-cell or single-nucleus approaches. Unsurprisingly, CGRP^PBN^ neurons showed high expression of *Calca*, along with other neuropeptide-related genes and pathways (**Figure 1D-E**). Interestingly, some of these neuropeptide targets have well-established links to appetite, consistent with the role that CGRP^PBN^ neurons play in appetitive behaviors, as discussed in the introduction.

In addition to producing a transcriptional profile of CGRP^PBN^ neurons, we also sought out to identify CGRP^PBN^ genes that were highly enriched in this population when compared to other neuron subtypes. To this end, we compared the CGRP^PBN^ neuron transcriptome to publicly available gene expression profiles of other neuronal subtypes across various brain regions.^39,41^ Importantly, while samples collected by our lab included equal numbers of male and female mice, and analyses were performed to detect sex differences between HA+ and HA- nuclei (**Supplementary Figure 2A-D**), the publicly available datasets used in this study included only male animals. As such, this analysis could not be performed across all cell types to determine whether highly enriched CGRP^PBN^ genes are sexually dimorphic. Nevertheless, our analysis found that, consistent with the previously published single-cell study,^11^ genes encoding the neuropeptides CGRP, galanin, neurotensin, and nesfatin-1 were significantly enriched in CGRP^PBN^ neurons (**Figure 2D**). These neuropeptides have established roles in appetite regulation and also exhibit significant overlap with reward-related brain regions.^19,52–55,57,59,60,72^ This is also in line with *in situ* hybridization studies, which mapped projections from CGRP^PBN^ neurons to various brain regions involved in reward in the mouse brain.^11^ Considering CGRP^PBN^ neurons are also enriched for *Oprm1* (**Supplementary Figure 3A**) along with other opioid-related genes such as *Penk* and *Oprl1* (**Figure 2D**), there may be a relationship between appetitive morphine consumption and CGRP^PBN^ neuron function. Further, given the recent interest in repurposing GLP-1 therapeutics for SUDs,^25,73–75^ the direct link between GLP-1 signaling and CGRP-mediated appetite regulation,^20,22,23^ and the current availability of FDA-approved CGRP receptor inhibitor therapeutics, understanding the role of CGRP^PBN^ neurons in OUD-related behaviors is of high translational relevance.

Consistent with our molecular findings suggesting that CGRP^PBN^ neurons may be sensitive to opioids, cFos immunofluorescence analysis revealed that CGRP^PBN^ neurons become activated by abstinence following chronic (5-day) morphine administration (**Figure 3B-C**). There is significant evidence that CGRP^PBN^ neurons responsive to aversive stimuli project to the CeA. ^13,60,76–78^ Importantly, previous studies have shown that reduced CGRP release decreases activation of the CeA, and when CGRP receptors in the CeA are inhibited, electrophysiological and behavioral responses to CGRP activation are attenuated. ^12–14,78^ This is particularly noteworthy in the context of this study because the CeA also has an established role in mediating opioid withdrawal, and its activity increases during this aversive experience.^79–83^ Given that our results on the timing of CGRP^PBN^ activation follow the same timeline for many acute withdrawal behaviors,^61,62^ we suggest that projections from CGRP^PBN^ neurons to the CeA may mediate some of the motivation- and affective-related behaviors characteristic of opioid withdrawal. However, behavioral assessments that measure both somatic and affective aspects of opioid withdrawal will be necessary to test this hypothesis.

In the present study, we first established that at a single dose of morphine, CNO-mediated chemogenetic inhibition of CGRP^PBN^ neurons reduced morphine intake in hM4Di- (**Figure 4D**) but not DIO-mCherry-expressing Calca^Cre+/-^ mice(**Figure 4C**). We then repeated this intervention within the context of a dose-response model to more accurately determine the role CGRP^PBN^ neurons may play in opioid reinforcement. In this experiment, we showed that there was a significant reduction in morphine intake at the lowest 2 of the 4 doses tested when CGRP^PBN^ neurons were chemogenetically inhibited (**Figure 5B**). These results suggest that CGRP^PBN^ neurons can regulate morphine reinforcement. Moreso, the direction of this regulation is consistent with previous studies showing that knockdown of the CGRP receptor in D2-expressing neurons of the NAc reduced oxycodone self-administration.^15^ In contrast, however, this same study found that when knockdown occurred in D1-expressing neurons, oxycodone self-administration instead increased. Further, studies implicating CGRP^PBN^ neurons in appetitive behaviors found that CGRP^PBN^ neuronal activation, rather than inhibition, induces anorexic effects via projections from the NTS to CGRP^PBN^ neurons and from CGRP^PBN^ neurons to the CeA.^19,20^ Contextualizing our data within this framework suggests that CGRP^PBN^ neurons may have projection-specific functions that have yet to be explored in the context of OUD. This would be consistent with the established roles of this neuronal population in processing multiple types of stimuli and is additionally supported by immunofluorescence studies showing that anatomical projections from *Calca-*expressing neurons follow at least 2 distinct pathways based on their gene expression profiles,^11^ which have not yet been functionally characterized. To determine the mechanisms by which CGRP^PBN^ neurons regulate morphine reinforcement and motivated behavior, studies using projection-specific activation and inhibition will be necessary.

Finally, we investigated the effect of CGRP^PBN^ neuronal inhibition on relapse-related behaviors. In this experiment, we found a significant increase in seeking behavior during acute abstinence (**Figure 5D**), which persisted after 21 days of forced home-cage abstinence (**Figure 5D**). However, chemogenetic inhibition of CGRP^PBN^ neurons during chronic abstinence did not alter seeking behavior, suggesting that the behavioral regulation of relapse may be independent of the activity of CGRP^PBN^ neurons. Notwithstanding, opioid relapse is highly associated with initial craving and severity of withdrawal,^84,85^ thus it is still possible that CGRP^PBN^ neurons may regulate initial levels of craving during the acute abstinence session, which we did not test. This is supported by the initial increase in seeking behavior compared to baseline at this time point (**Figure 5E**), which corresponded to a significantly elevated level of CGRP^PBN^ activity in our cFos staining data (**Figure 3B-C**). Inhibition of CGRP^PBN^ neurons at this early time point (acute abstinence) may better illustrate the relationship between CGRP^PBN^ neuronal activity and both withdrawal-related craving and future relapse risk.

## Conclusions

In this study, we employed a viral nuclear-labeling approach to isolate a pure population of CGRP^PBN^ neurons to generate a cell-specific CGRP^PBN^ transcriptome. Comparison of the CGRP^PBN^ transcriptome to those of neuronal and non-neuronal cell types across different brain regions revealed CGRP^PBN^ enrichment of genes associated with pain, stress, neuropeptide signaling, and inhibitory synapses. Interestingly, the neuropeptides most highly enriched in CGRP^PBN^ neurons are significantly implicated in regulating behaviors associated with appetite and reward. We also show that morphine abstinence disinhibits CGRP^PBN^ neuron activity, and that DREADD-mediated inhibition of this population reduces morphine intake. While further work is necessary to fully understand the mechanisms underlying these behavioral effects, the association between CGRP^PBN^ signaling, appetite, and reward strongly suggests that these neurons are involved in regulating the motivational aspects of morphine-taking. Currently, CGRP receptor inhibitors are FDA-approved for the treatment and prevention of migraines. Given this commercial availability and lack of µOR engagement, the translational impact of developing this target could be significant, with immediate implications for the treatment of OUD.

## Acknowledgments

We would like to thank Dr. Richard Palmiter (University of Washington; HHMI) for generously sharing with us a founding pair of Calca^tm1.1(Cre/EGFP)Rpa^ mice, as well as providing technical advice on genotyping.

## Funding

DP1DA051828 (LMT)

F30DA060666 (LLB)

Kind gift from the Shipley Foundation (LMT)

## Author contributions

Conceptualization: LLB, LMT

Methodology: LLB, AVM, LMT

Software: LLB, AVM

Investigation: LLB, AVM, NMK, EPM, FMBG, LMT

Visualization: LLB

Data Curation: LLB, LMT

Supervision: FMBG, LMT

Writing—original draft: LLB, LMT

Writing—review & editing: LLB, AVM, FMBG, LMT

## Data and materials availability

All next-generation sequencing files associated with this study, as well as the code that was used to pre-process and run differential expression, are available online at https://github.com/avm27/CGRPReinforcement. All data files used in processing and analysis will be available from the GEO/SRA upon publication and/or upon request.

## References

1. Ahmad F, Cisewski J, Rossen L, Sutton P. Provisional drug overdose death counts. https://www.cdc.gov/nchs/nvss/vsrr/drug-overdose-data.htm

2. CDC. Understanding the Opioid Overdose Epidemic. Accessed July 19 2023, https://www.cdc.gov/opioids/basics/epidemic.html

3. DEA. 2019 National Drug Threat Assessment. U.S. Department of Justice.

4. Larochelle MR, Bernson D, Land T, et al. Medication for Opioid Use Disorder After Nonfatal Opioid Overdose and Association With Mortality: A Cohort Study. Ann Intern Med. Aug 7 2018;169(3):137–145. doi:10.7326/m17-3107

5. NIDA, NIH, HHS. 2023. https://nida.nih.gov/research-topics/opioids/medications-opioid-overdose-withdrawal-addiction-infographic

6. Morgan JR, Schackman BR, Leff JA, Linas BP, Walley AY. Injectable naltrexone, oral naltrexone, and buprenorphine utilization and discontinuation among individuals treated for opioid use disorder in a United States commercially insured population. J Subst Abuse Treat. Feb 2018;85:90–96. doi:10.1016/j.jsat.2017.07.001

7. Jiang X, Guy GP, Jr., Dever JA, et al. Association Between Length of Buprenorphine or Methadone Use and Nonprescribed Opioid Use Among Individuals with Opioid Use Disorder: A Cohort Study. Subst Use Addctn J. Apr 2025;46(2):266–279. doi:10.1177/29767342241266038

8. Greenwald MK, Sogbesan T, Moses TEH. Relationship between opioid cross-tolerance during buprenorphine stabilization and return to opioid use during buprenorphine dose tapering. Psychopharmacology (Berl). Jun 2024;241(6):1151–1160. doi:10.1007/s00213-024-06549-1

9. Volkow ND, Blanco C. Medications for opioid use disorders: clinical and pharmacological considerations. The Journal of Clinical Investigation. 11/25/ 2019;130(1):10–13. doi:10.1172/JCI134708

10. Alhadeff AL, Holland RA, Zheng H, Rinaman L, Grill HJ, De Jonghe BC. Excitatory Hindbrain-Forebrain Communication Is Required for Cisplatin-Induced Anorexia and Weight Loss. J Neurosci. Jan 11 2017;37(2):362–370. doi:10.1523/jneurosci.2714-16.2016

11. Pauli JL, Chen JY, Basiri ML, et al. Molecular and anatomical characterization of parabrachial neurons and their axonal projections. eLife. 2022/11/01 2022;11:e81868. doi:10.7554/eLife.81868

12. Torres-Rodriguez JM, Wilson TD, Singh S, et al. The parabrachial to central amygdala pathway is critical to injury-induced pain sensitization in mice. Neuropsychopharmacology. Feb 2024;49(3):508–520. doi:10.1038/s41386-023-01673-6

13. Palmiter RD. The Parabrachial Nucleus: CGRP Neurons Function as a General Alarm. Trends Neurosci. May 2018;41(5):280–293. doi:10.1016/j.tins.2018.03.007

14. Okutsu Y, Takahashi Y, Nagase M, Shinohara K, Ikeda R, Kato F. Potentiation of NMDA receptor-mediated synaptic transmission at the parabrachial-central amygdala synapses by CGRP in mice. Mol Pain. Jan-Dec 2017;13:1744806917709201. doi:10.1177/1744806917709201

15. Zhang Y, Ben Nathan J, Moreno A, et al. Calcitonin receptor signaling in nucleus accumbens D1R- and D2R-expressing medium spiny neurons bidirectionally alters opioid taking in male rats. Neuropsychopharmacology. 2023/12/01 2023;48(13):1878–1888. doi:10.1038/s41386-023-01634-z

16. Powell R, Young VA, Pryce KD, et al. Inhibiting endocytosis in CGRP+ nociceptors attenuates inflammatory pain-like behavior. Nature Communications. 2021/10/04 2021;12(1):5812. doi:10.1038/s41467-021-26100-6

17. Fisher LA, Kikkawa DO, Rivier JE, et al. Stimulation of noradrenergic sympathetic outflow by calcitonin gene-related peptide. Nature. Oct 6-12 1983;305(5934):534–6. doi:10.1038/305534a0

18. Michael CC, Anna B, Lindsey AS, Domenico T, Olivia U, Mary MH. Parabrachial Complex: A Hub for Pain and Aversion. The Journal of Neuroscience. 2019;39(42):8225. doi:10.1523/JNEUROSCI.1162-19.2019

19. Carter ME, Soden ME, Zweifel LS, Palmiter RD. Genetic identification of a neural circuit that suppresses appetite. Nature. Nov 7 2013;503(7474):111–4. doi:10.1038/nature12596

20. Roman CW, Derkach VA, Palmiter RD. Genetically and functionally defined NTS to PBN brain circuits mediating anorexia. Nat Commun. Jun 15 2016;7:11905. doi:10.1038/ncomms11905

21. Campos CA, Bowen AJ, Schwartz MW, Palmiter RD. Parabrachial CGRP Neurons Control Meal Termination. Cell Metab. May 10 2016;23(5):811–20. doi:10.1016/j.cmet.2016.04.006

22. Richard JE, Farkas I, Anesten F, et al. GLP-1 receptor stimulation of the lateral parabrachial nucleus reduces food intake: neuroanatomical, electrophysiological, and behavioral evidence. Endocrinology. Nov 2014;155(11):4356–67. doi:10.1210/en.2014-1248

23. Swick JC, Alhadeff AL, Grill HJ, et al. Parabrachial Nucleus Contributions to Glucagon-Like Peptide-1 Receptor Agonist-Induced Hypophagia. Neuropsychopharmacology. Jul 2015;40(8):2001–14. doi:10.1038/npp.2015.50

24. Hayes MR, Bradley L, Grill HJ. Endogenous hindbrain glucagon-like peptide-1 receptor activation contributes to the control of food intake by mediating gastric satiation signaling. Endocrinology. Jun 2009;150(6):2654–9. doi:10.1210/en.2008-1479

25. Tuesta LM, Chen Z, Duncan A, et al. GLP-1 acts on habenular avoidance circuits to control nicotine intake. Nat Neurosci. May 2017;20(5):708–716. doi:10.1038/nn.4540

26. Zhang Y, Kahng MW, Elkind JA, et al. Activation of GLP-1 receptors attenuates oxycodone taking and seeking without compromising the antinociceptive effects of oxycodone in rats. Neuropsychopharmacology. Feb 2020;45(3):451–461. doi:10.1038/s41386-019-0531-4

27. Urbanik LA, Acharya NK, Grigson PS. Acute treatment with the glucagon-like peptide-1 receptor agonist, liraglutide, reduces cue- and drug-induced fentanyl seeking in rats. Brain Res Bull. Oct 15 2022;189:155–162. doi:10.1016/j.brainresbull.2022.08.023

28. Farokhnia M, Tazare J, Pince CL, et al. Glucagon-like peptide-1 receptor agonists, but not dipeptidyl peptidase-4 inhibitors, reduce alcohol intake. J Clin Invest. May 1 2025;135(9)doi:10.1172/jci188314

29. Douton JE, Horvath N, Mills-Huffnagle S, Nyland JE, Hajnal A, Grigson PS. Glucagon-like peptide-1 receptor agonist, liraglutide, reduces heroin self-administration and drug-induced reinstatement of heroin-seeking behaviour in rats. Addict Biol. Mar 2022;27(2):e13117. doi:10.1111/adb.13117

30. Furdui A, da Silveira Scarpellini C, Montandon G. Anatomical distribution of µ-opioid receptors, neurokinin-1 receptors, and vesicular glutamate transporter 2 in the mouse brainstem respiratory network. J Neurophysiol. Jul 1 2024;132(1):108–129. doi:10.1152/jn.00478.2023

31. Zhang XY, Li Q, Dong Y, et al. Mu-Opioid Receptors Expressed in Glutamatergic Neurons are Essential for Morphine Withdrawal. Neurosci Bull. Oct 2020;36(10):1095–1106. doi:10.1007/s12264-020-00515-5

32. Allen Reference Atlases: Atlas Viewer. https://atlas.brain-map.org

33. Simon MJ, Molina F, Puerto A. Conditioned place preference but not rewarding self-stimulation after electrical activation of the external lateral parabrachial nucleus. Behav Brain Res. Dec 28 2009;205(2):443–9. doi:10.1016/j.bbr.2009.07.028

34. Simon MJ, Garcia R, Zafra MA, Molina F, Puerto A. Learned preferences induced by electrical stimulation of a food-related area of the parabrachial complex: effects of naloxone. Neurobiol Learn Mem. Mar 2007;87(3):332–42. doi:10.1016/j.nlm.2006.09.009

35. Simon MJ, Garcia R, Puerto A. Concurrent stimulation-induced place preference in lateral hypothalamus and parabrachial complex: differential effects of naloxone. Behav Brain Res. Nov 20 2011;225(1):311–6. doi:10.1016/j.bbr.2011.07.029

36. Hurtado MM, García R, Puerto A. Naloxone blocks the aversive effects of electrical stimulation of the parabrachial complex in a place discrimination task. Neurobiol Learn Mem. Dec 2016;136:21–27. doi:10.1016/j.nlm.2016.09.011

37. Mussetto V, Teuchmann HL, Heinke B, et al. Opioids Induce Bidirectional Synaptic Plasticity in a Brainstem Pain Center in the Rat. The Journal of Pain. 2023/09/01/ 2023;24(9):1664–1680. 10.1016/j.jpain.2023.05.001

38. Salmon AM, Damaj MI, Marubio LM, Epping-Jordan MP, Merlo-Pich E, Changeux JP. Altered neuroadaptation in opiate dependence and neurogenic inflammatory nociception in alpha CGRP-deficient mice. Nat Neurosci. Apr 2001;4(4):357–8. doi:10.1038/86001

39. Tuesta LM, Djekidel MN, Chen R, et al. In vivo nuclear capture and molecular profiling identifies Gmeb1 as a transcriptional regulator essential for dopamine neuron function. Nat Commun. Jun 7 2019;10(1):2508. doi:10.1038/s41467-019-10267-0

40. Krashes MJ, Koda S, Ye C, et al. Rapid, reversible activation of AgRP neurons drives feeding behavior in mice. J Clin Invest. Apr 2011;121(4):1424–8. doi:10.1172/jci46229

41. Mo A, Mukamel EA, Davis FP, et al. Epigenomic Signatures of Neuronal Diversity in the Mammalian Brain. Neuron. Jun 17 2015;86(6):1369–84. doi:10.1016/j.neuron.2015.05.018

42. Felix Krueger F. J., PE, Ebrahim Afyounian, Michael Weinstein, Benjamin Schuster-Boeckler, Gert Hulselmans, & sclamons. TrimGalore: v0.6.10 - add default decompression path (0.6.10). Zenodo. 2023;

43. M. M. Cutadapt removes adapter sequences from high-throughput sequencing reads. EMBnetjournal. 2011;17(1):10–12.

44. Dobin A, Davis CA, Schlesinger F, et al. STAR: ultrafast universal RNA-seq aligner. Bioinformatics. Jan 1 2013;29(1):15–21. doi:10.1093/bioinformatics/bts635

45. Pertea M, Pertea GM, Antonescu CM, Chang TC, Mendell JT, Salzberg SL. StringTie enables improved reconstruction of a transcriptome from RNA-seq reads. Nat Biotechnol. Mar 2015;33(3):290–5. doi:10.1038/nbt.3122

46. Love MI, Huber W, Anders S. Moderated estimation of fold change and dispersion for RNA-seq data with DESeq2. Genome Biol. 2014;15(12):550. doi:10.1186/s13059-014-0550-8

47. Wu T, Hu E, Xu S, et al. clusterProfiler 4.0: A universal enrichment tool for interpreting omics data. Innovation (Camb). Aug 28 2021;2(3):100141. doi:10.1016/j.xinn.2021.100141

48. Yu G, Wang LG, Han Y, He QY. clusterProfiler: an R package for comparing biological themes among gene clusters. Omics. May 2012;16(5):284–7. doi:10.1089/omi.2011.0118

49. Wickham H. Ggplot2: elegant graphics for data analysis. Springer,. 2009;

50. Schindelin J, Arganda-Carreras I, Frise E, et al. Fiji: an open-source platform for biological-image analysis. Nat Methods. Jun 28 2012;9(7):676–82. doi:10.1038/nmeth.2019

51. Vilca SJ, Margetts AV, Höglund L, et al. Microglia contribute to methamphetamine reinforcement and reflect persistent transcriptional and morphological adaptations to the drug. Brain Behav Immun. Aug 2024;120:339–351. doi:10.1016/j.bbi.2024.05.038

52. Qualls-Creekmore E, Yu S, Francois M, et al. Galanin-Expressing GABA Neurons in the Lateral Hypothalamus Modulate Food Reward and Noncompulsive Locomotion. J Neurosci. Jun 21 2017;37(25):6053–6065. doi:10.1523/jneurosci.0155-17.2017

53. Scheller KJ, Williams SJ, Lawrence AJ, Djouma E. The galanin-3 receptor antagonist, SNAP 37889, suppresses alcohol drinking and morphine self-administration in mice. Neuropharmacology. May 15 2017;118:1–12. doi:10.1016/j.neuropharm.2017.03.004

54. Brady LS, Smith MA, Gold PW, Herkenham M. Altered expression of hypothalamic neuropeptide mRNAs in food-restricted and food-deprived rats. Neuroendocrinology. Nov 1990;52(5):441–7. doi:10.1159/000125626

55. Ratner C, Skov LJ, Raida Z, et al. Effects of Peripheral Neurotensin on Appetite Regulation and Its Role in Gastric Bypass Surgery. Endocrinology. Sep 2016;157(9):3482–92. doi:10.1210/en.2016-1329

56. Khan R, Laumet G, Leinninger GM. Hungry for relief: Potential for neurotensin to address comorbid obesity and pain. Appetite. Sep 1 2024;200:107540. doi:10.1016/j.appet.2024.107540

57. Javed A, Kamradt MC, Van de Kar LD, Gray TS. D-Fenfluramine induces serotonin-mediated Fos expression in corticotropin-releasing factor and oxytocin neurons of the hypothalamus, and serotonin-independent Fos expression in enkephalin and neurotensin neurons of the amygdala. Neuroscience. Mar 1999;90(3):851–8. doi:10.1016/s0306-4522(98)00523-5

58. Bayless DW, Davis CO, Yang R, et al. A neural circuit for male sexual behavior and reward. Cell. Aug 31 2023;186(18):3862–3881.e28. doi:10.1016/j.cell.2023.07.021

59. Greenwell TN, Martin-Schild S, Inglis FM, Zadina JE. Colocalization and shared distribution of endomorphins with substance P, calcitonin gene-related peptide, gamma-aminobutyric acid, and the mu opioid receptor. J Comp Neurol. Jul 10 2007;503(2):319–33. doi:10.1002/cne.21374

60. Campos CA, Bowen AJ, Roman CW, Palmiter RD. Encoding of danger by parabrachial CGRP neurons. Nature. Mar 29 2018;555(7698):617–622. doi:10.1038/nature25511

61. Brewer AL, Lewis CC, Eggerman L, et al. Modeling spontaneous opioid withdrawal in male and female outbred mice using traditional endpoints and hyperalgesia. Behav Pharmacol. Apr 1 2023;34(2-3):112–122. doi:10.1097/fbp.0000000000000714

62. Ozdemir D, Allain F, Kieffer BL, Darcq E. Advances in the characterization of negative affect caused by acute and protracted opioid withdrawal using animal models. Neuropharmacology. Jul 1 2023;232:109524. doi:10.1016/j.neuropharm.2023.109524

63. Nett KE, LaLumiere RT. Pair housing does not alter incubation of craving, extinction, and reinstatement after heroin self-administration in female and male rats. Behav Neurosci. Apr 2023;137(2):111–119. doi:10.1037/bne0000544

64. Bossert JM, Caldwell KE, Korah H, et al. Effect of chronic delivery of the NOP/MOR partial agonist AT-201 and NOP antagonist J-113397 on heroin relapse in a rat model of opioid maintenance. Psychopharmacology (Berl). Dec 2024;241(12):2497–2511. doi:10.1007/s00213-024-06678-7

65. Huerta Sanchez LL, Sankaran M, Li TL, et al. Profiling prefrontal cortex protein expression in rats exhibiting an incubation of cocaine craving following short-access self-administration procedures. Front Psychiatry. 2022;13:1031585. doi:10.3389/fpsyt.2022.1031585

66. Bentzley BS, Barth KS, Back SE, Aronson G, Book SW. Patient Perspectives Associated with Intended Duration of Buprenorphine Maintenance Therapy. J Subst Abuse Treat. Sep 2015;56:48–53. doi:10.1016/j.jsat.2015.04.002

67. Wyse JJ, Eckhardt A, Waller D, et al. Patients’ Perspectives on Discontinuing Buprenorphine for the Treatment of Opioid Use Disorder. J Addict Med. May-Jun 01 2024;18(3):300–305. doi:10.1097/adm.0000000000001292

68. Sadek J, Saunders J. Treatment retention in opioid agonist therapy: comparison of methadone versus buprenorphine/naloxone by analysis of daily-witnessed dispensed medication in a Canadian Province. BMC Psychiatry. Jul 30 2022;22(1):516. doi:10.1186/s12888-022-04175-9

69. Ahmed FZ, Andraka-Christou B, Clark MH, Totaram R, Atkins DN, del Pozo B. Barriers to medications for opioid use disorder in the court system: provider availability, provider “trustworthiness,” and cost. Health & Justice. 2022/07/27 2022;10(1):24. doi:10.1186/s40352-022-00188-4

70. Jones CM, Han B, Baldwin GT, Einstein EB, Compton WM. Use of Medication for Opioid Use Disorder Among Adults With Past-Year Opioid Use Disorder in the US, 2021. JAMA Network Open. 2023;6(8):e2327488–e2327488. doi:10.1001/jamanetworkopen.2023.27488

71. Pollock TA, Margetts AV, Vilca SJ, Tuesta LM. Cocaine taking and craving produce distinct transcriptional profiles in dopamine neurons. bioRxiv. Oct 12 2024;doi:10.1101/2024.10.11.617923

72. Dore R, Krotenko R, Reising JP, et al. Nesfatin-1 decreases the motivational and rewarding value of food. Neuropsychopharmacology. Sep 2020;45(10):1645–1655. doi:10.1038/s41386-020-0682-3

73. Eren-Yazicioglu CY, Yigit A, Dogruoz RE, Yapici-Eser H. Can GLP-1 Be a Target for Reward System Related Disorders? A Qualitative Synthesis and Systematic Review Analysis of Studies on Palatable Food, Drugs of Abuse, and Alcohol. Front Behav Neurosci. 2020;14:614884. doi:10.3389/fnbeh.2020.614884

74. Schmidt HD, Mietlicki-Baase EG, Ige KY, et al. Glucagon-Like Peptide-1 Receptor Activation in the Ventral Tegmental Area Decreases the Reinforcing Efficacy of Cocaine. Neuropsychopharmacology. Jun 2016;41(7):1917–28. doi:10.1038/npp.2015.362

75. Shevchouk OT, Tufvesson-Alm M, Jerlhag E. An Overview of Appetite-Regulatory Peptides in Addiction Processes; From Bench to Bed Side. Front Neurosci. 2021;15:774050. doi:10.3389/fnins.2021.774050

76. Kocorowski LH, Helmstetter FJ. Calcitonin gene-related peptide released within the amygdala is involved in Pavlovian auditory fear conditioning. Neurobiol Learn Mem. Mar 2001;75(2):149–63. doi:10.1006/nlme.2000.3963

77. Missig G, Mei L, Vizzard MA, et al. Parabrachial Pituitary Adenylate Cyclase-Activating Polypeptide Activation of Amygdala Endosomal Extracellular Signal-Regulated Kinase Signaling Regulates the Emotional Component of Pain. Biol Psychiatry. Apr 15 2017;81(8):671–682. doi:10.1016/j.biopsych.2016.08.025

78. Han S, Soleiman MT, Soden ME, Zweifel LS, Palmiter RD. Elucidating an Affective Pain Circuit that Creates a Threat Memory. Cell. Jul 16 2015;162(2):363–374. doi:10.1016/j.cell.2015.05.057

79. Wooldridge LM, Wu JWK, Jo AY, et al. Central amygdalar PKCδ neurons mediate fentanyl withdrawal. bioRxiv. 2025:2025.02.21.639538. doi:10.1101/2025.02.21.639538

80. Wang L, Shen M, Jiang C, Ma L, Wang F. Parvalbumin Interneurons of Central Amygdala Regulate the Negative Affective States and the Expression of Corticotrophin-Releasing Hormone During Morphine Withdrawal. International Journal of Neuropsychopharmacology. 2016;19(11)doi:10.1093/ijnp/pyw060

81. Li YQ, Li FQ, Wang XY, et al. Central amygdala extracellular signal-regulated kinase signaling pathway is critical to incubation of opiate craving. J Neurosci. Dec 3 2008;28(49):13248–57. doi:10.1523/jneurosci.3027-08.2008

82. Frenois F, Stinus L, Blasi FD, Cador M, Moine CL. A Specific Limbic Circuit Underlies Opiate Withdrawal Memories. The Journal of Neuroscience. 2005;25(6):1366–1374. doi:10.1523/jneurosci.3090-04.2005

83. Bajo M, Madamba SG, Roberto M, Siggins GR. Acute morphine alters GABAergic transmission in the central amygdala during naloxone-precipitated morphine withdrawal: role of cyclic AMP. Original Research. Frontiers in Integrative Neuroscience. 2014-June-04 2014;8 doi:10.3389/fnint.2014.00045

84. Tsui JI, Anderson BJ, Strong DR, Stein MD. Craving predicts opioid use in opioid-dependent patients initiating buprenorphine treatment: a longitudinal study. Am J Drug Alcohol Abuse. Mar 2014;40(2):163–9. doi:10.3109/00952990.2013.848875

85. Vafaie N, Kober H. Association of Drug Cues and Craving With Drug Use and Relapse: A Systematic Review and Meta-analysis. JAMA Psychiatry. 2022;79(7):641–650. doi:10.1001/jamapsychiatry.2022.1240

