## Supplementary Figures for "Parabrachial CGRP Neurons Regulate Opioid Reinforcement"

**Supplementary Material**

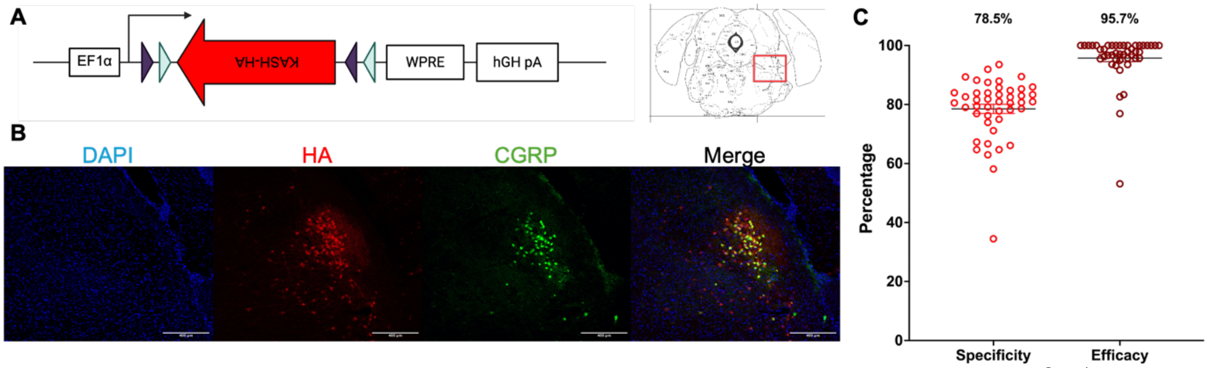


**Supplementary Figure 1: Validation of KASH-HA.** Male and Female Calca$^{Cre+/-}$ mice were bilaterally injected with an AAV-DIO-KASH-HA virus. Two weeks later, the parabrachial nucleus was dissected, and nuclei were isolated using Fluorescence Assisted Nuclear Sorting (FANS). These nuclei were then used for RNAseq analysis. **A)** Diagram of the DIO-KASH-HA vector adapted from Tuesta et. al. (2019). **B)** Representative micrographs showing infection with HA virus (red) co-localized to Cre+ CGRP neurons (green). Scale bar: 400 µm. Schematic Image from Allen Brain Atlas representing region images were taken from shown just above images **C)** Quantification of infection efficiency of DIO-KASH-HA construct. N=6, with 2-12 sections per animal.

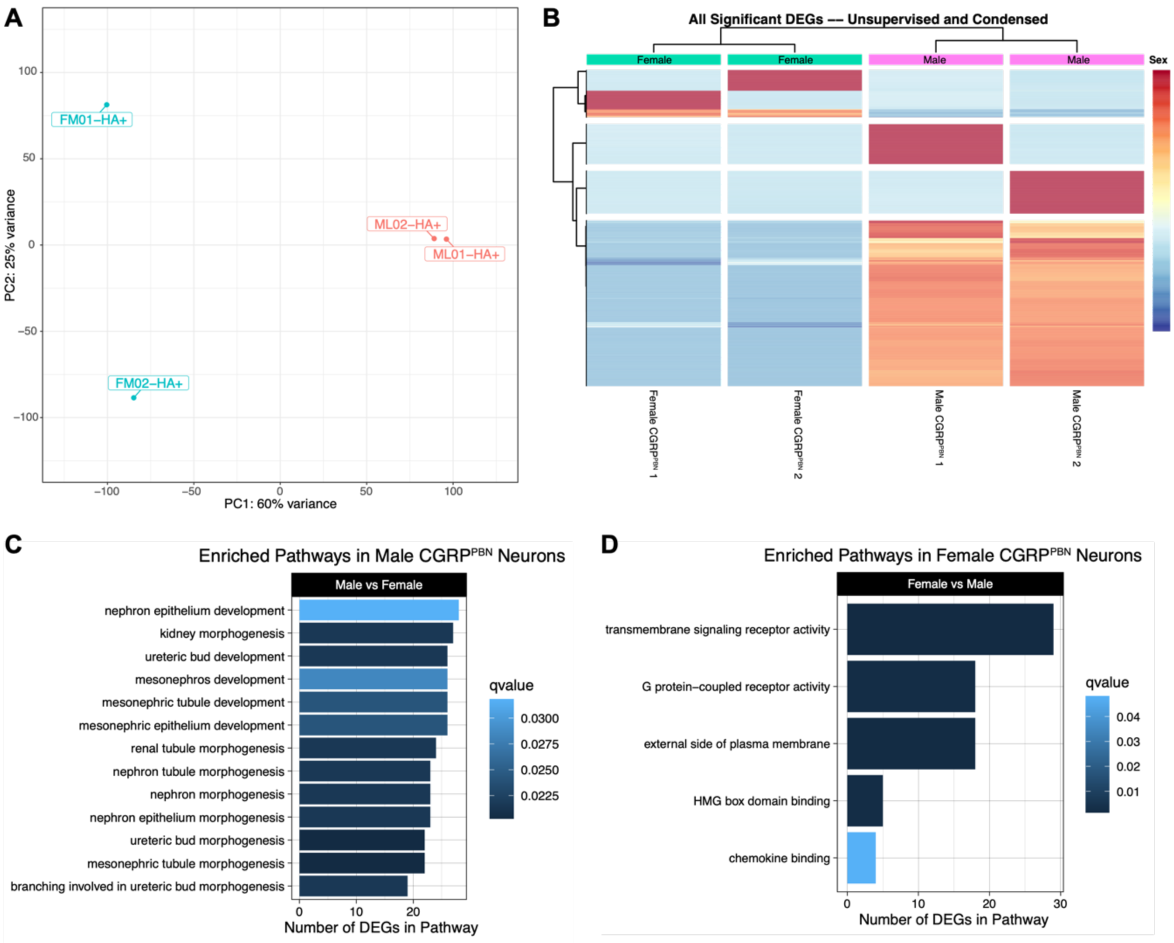


**Supplementary Figure 2: CGRP$^{PBN}$ Neurons are Sexually Dimorphic.** Male and Female Calca$^{Cre+/-}$ mice were bilaterally injected with an AAV-DIO-KASH-HA virus. Two weeks later, the parabrachial nucleus was dissected, and nuclei were isolated using Fluorescence Assisted Nuclear Sorting (FANS). These nuclei were then used for RNA-seq analysis. **A)** Principal Component Analysis of Variance between male (salmon) and female (cyan) CGRP$^{PBN}$ neuronal nuclei. **B)** Unsupervised clustering heatmap of normalized counts (rlog transformed) of all significant differentially expressed genes (*padj* < 0.05) between male and female CGRP$^{PBN}$ neuronal nuclei. **C)** Significantly enriched Gene Ontology (GO) pathways in male compared to female CGRP$^{PBN}$ neuronal nuclei ($padj_{genes}$ < 0.05, $padj_{pathways}$ < 0.05, $q\text{-}value_{pathways}$ < 0.05, L2FC > 1, Fold Enrichment ≥ 2). **D)** Significantly enriched Gene Ontology (GO) pathways in female compared to male CGRP$^{PBN}$ neuronal nuclei ($padj_{genes}$ < 0.05, $padj_{pathways}$ < 0.05, $q\text{-}value_{pathways}$ < 0.05, L2FC > 1, Fold Enrichment ≥ 2).

**A**

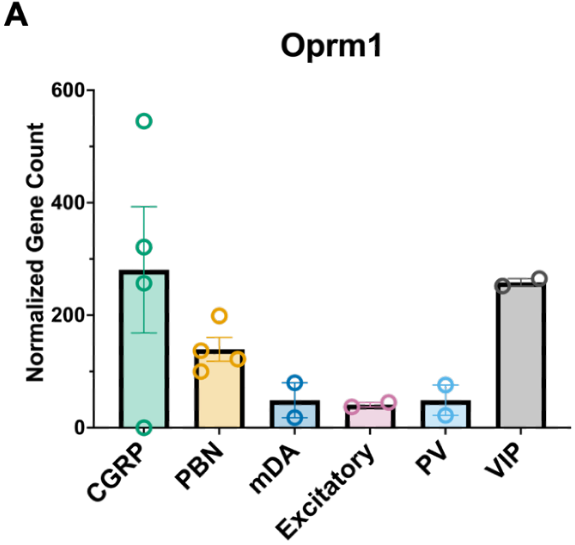


**Supplementary Figure 3: *Oprm1* is highly expressed in CGRP$^{PBN}$ neurons.** Gene expression data from publicly available datasets (GEO: GSE106956 and GSE63137) were processed alongside CGRP$^{PBN}$ and non-CGRP PBN samples and underwent differential gene expression and functional enrichment analyses. **A)** Normalized gene expression plot for *Oprm1.* n=2-4 per cell type; Error bars represent mean ± SEM.

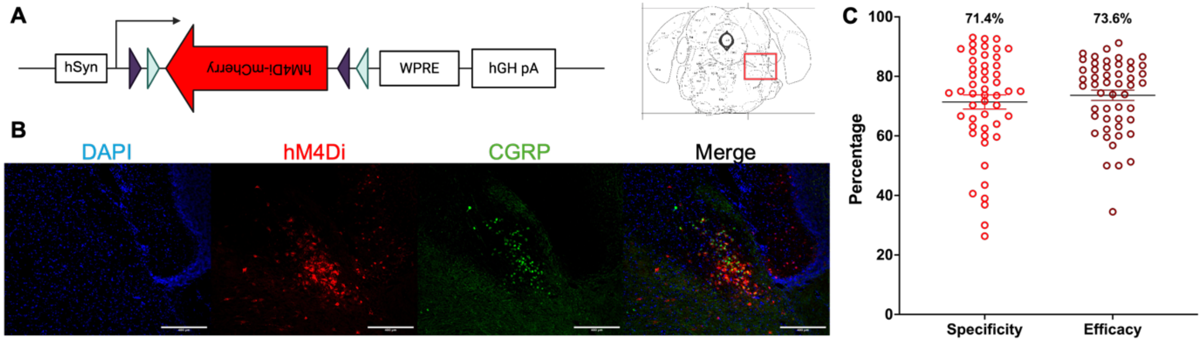


**Supplementary Figure 4: Validation of inhibitory DREADD (hM4Di) virus.** Male and female Calca$^{Cre+/-}$ mice were injected with an inhibitory DREADD or control virus in bilateral PBN. Two weeks later, mice underwent jugular catheterization and were trained to self-administer morphine during daily 2-hour sessions at FR3TO20. CNO or vehicle was administered 30 minutes prior to behavior session. During CNO test sessions, mice received either CNO (7.5 mg/kg, I.P) or vehicle 30 minutes prior to session start. **A)** Diagram of the DIO-hM4Di vector. **B)** Representative micrographs showing infection with hM4Di virus (red) co-localized to Cre+ CGRP neurons (green). Scale bar: 400 μm. Schematic Image from Allen Brain Atlas representing region images were taken from shown just above images **C)** Quantification of infection efficiency of DIO-hM4Di construct. N=4, with 11-15 sections per animal. Error bars represent mean ± SEM.

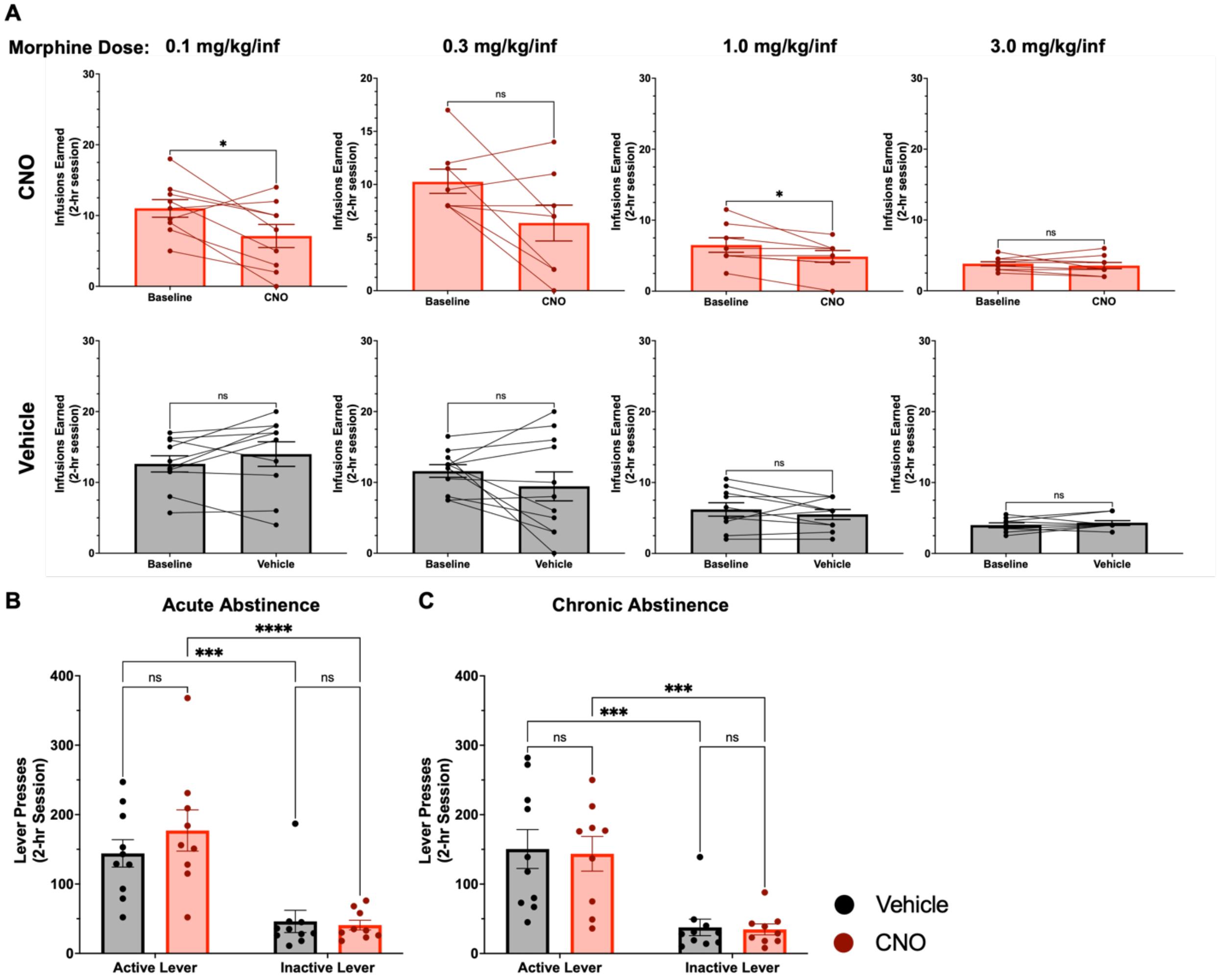


**Supplementary Figure 5: CGRP$^{PBN}$ neurons regulate morphine reinforcement, but not morphine-seeking.** Male and Female Calca$^{Cre+/-}$ mice were injected with an inhibitory DREADD virus. Two weeks later, mice were implanted with jugular catheters and underwent operant training to respond for morphine intravenously in a dose-response model. During Chronic Abstinence, mice received either CNO (7.5 mg/kg, I.P) or vehicle 30 minutes prior to session start. **A)** Morphine infusions earned by vehicle- and CNO-administered animals in baseline and test sessions. Doses are listed above each column in mg/kg/infusion (0.1: Paired t-test, $p=0.0345$; 1.0: Paired t-test, $p=0.0354$). **B)** Number of total Active or Inactive Lever Presses during Acute Abstinence. Two-way ANOVA- Effect of Lever ($F(1,34)=35.34$, $p<0.0001$). **C)** Number of total Active or Inactive Lever Presses during Chronic Abstinence. Two-way ANOVA- Effect of Lever ($F(1,34)=29.89$, $p<0.0001$). **A-C)** *$p<0.05$,***$p<0.001$, ****$p<0.0001$; Error bars represent mean ± SEM. N=9-11 animals per group per comparison. Vehicle is represented in black, while CNO-treated animals are represented in red.
